# Sequence-dependent mechanics of collagen reflect its structural and functional organization

**DOI:** 10.1101/2020.09.27.315929

**Authors:** Alaa Al-Shaer, Aaron Lyons, Yoshihiro Ishikawa, Billy G. Hudson, Sergei P. Boudko, Nancy R. Forde

## Abstract

Extracellular matrix mechanics influence diverse cellular functions, yet surprisingly little is known about the mechanical properties of their constituent collagen proteins. In particular, network-forming collagen IV, an integral component of basement membranes, has been far less studied than fibril-forming collagens. A key feature of collagen IV is the presence of interruptions in the triple-helix-defining (Gly-X-Y) sequence along its collagenous domain. Here, we used atomic force microscopy (AFM) to determine the impact of sequence heterogeneity on the local flexibility of collagen IV and of the fibril-forming collagen III. Our extracted flexibility profile of collagen IV reveals that it possesses highly heterogeneous mechanics, ranging from semi-flexible regions as found for fibril-forming collagens to a lengthy region of high flexibility towards its N terminus. A simple model in which flexibility is dictated only by the presence of interruptions fit the extracted profile reasonably well, providing insight into the alignment of chains and demonstrating that interruptions – particularly when coinciding in multiple chains – significantly enhance local flexibility. To a lesser extent, sequence variations within the triple helix lead to variable flexibility, as seen along the continuously triple-helical collagen III. We found this fibril-forming collagen to possess a high-flexibility region around its matrix-metalloprotease (MMP) binding site, suggesting a unique mechanical fingerprint of this region that is key for matrix remodeling. Surprisingly, proline content did not correlate with local flexibility in either collagen type. We also found that physiologically relevant changes in pH and chloride concentration did not alter the flexibility of collagen IV, indicating such environmental changes are unlikely to control its compaction during secretion. Although extracellular chloride ions play a role in triggering collagen IV network formation, they do not appear to modulate the structure of its collagenous domain.

**Significance Statement:** Collagens are the predominant proteins in vertebrates, forming diverse hierarchical structures to support cells and form connective tissues. Despite their mechanical importance, surprisingly little is established about the molecular encoding of mechanics. Here, we image single collagen proteins and find that they exhibit variable flexibility along their backbones. By comparing collagens with continuous and discontinuous triple-helix-forming sequences, we find that the type of helix interruption correlates with local flexibility, providing the first steps towards a much-needed map between sequence, structure, and mechanics in these large proteins. Our results inform our understanding of collagen’s ability to adopt compact conformations during cellular secretion and suggest a physical mechanism by which higher-order structure may be regulated by the distinct molecular properties of different collagens.

## Introduction

Collagen is the most abundant protein in the animal kingdom and represents one third of the total protein in the human body (1, 2). It is a major structural component of the extracellular matrix, contributing to the mechanical stability, organization, and shape of a wide variety of tissues. Twenty-eight distinct collagen types have been reported in humans, with different higher-order organizational structures (1–3). The most prevalent are fibril-forming collagens: these have a unique hierarchical structure whereby collagen molecules assemble in a parallel, staggered fashion into long fibrillar nanostructures which form higher-order structures (Fig. 1). These fibers are the predominant load- and tension-bearing structures in connective tissues. Network-forming collagens, such as collagen IV, associate end-on to form sheet-like networks rather than fibrils (Fig. 1) (4, 5). These collagen IV networks played an integral role in the evolution of multicellular life (3, 6). Collagen IV networks are predominantly found in basement membranes (BMs) where they provide mechanical support and anchorage for cells and tissues, and serve as a filtration barrier to macromolecules in organs such as the kidney (5, 7). The mechanics of fibril-forming collagens have been the subject of numerous studies at the fibrillar and – to a lesser extent – molecular levels (8–10, and references therein). By contrast, the mechanics of network-forming collagens are far less studied (11). Similarly, knowledge of the structure of collagen IV lags significantly behind fibril-forming collagens, at both the supramolecular and molecular scales.

**Fig. 1.**
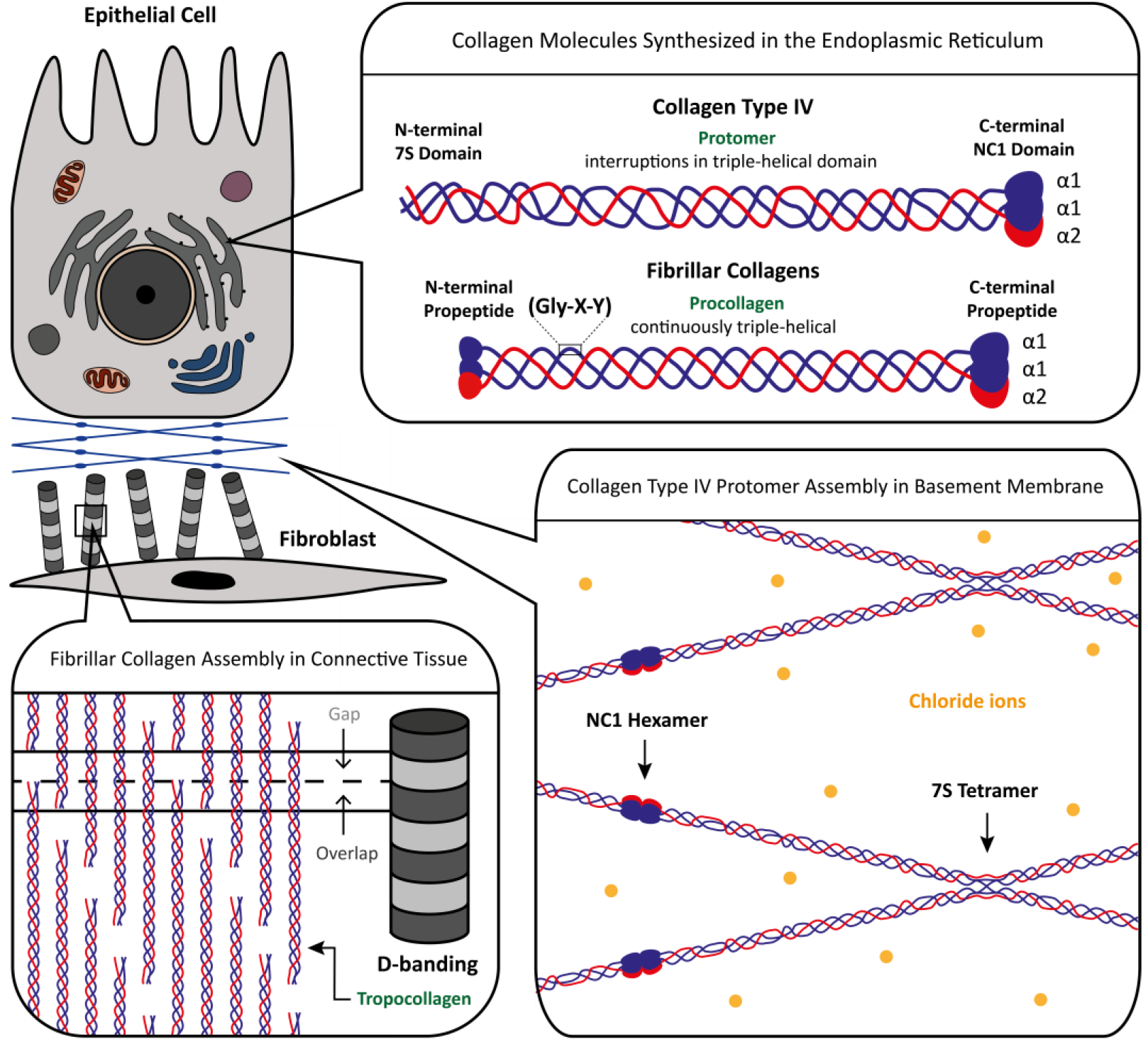
Localization and supramolecular structures of fibril-forming and network-forming collagens. All collagen chains are synthesized in the endoplasmic reticulum. The folded proteins are secreted from the cell, where they can form a variety of supramolecular structures in different tissues. The fibril-forming collagen molecule, procollagen, is post-translationally processed whereby its propeptides are cleaved off to yield tropocollagen. Tropocollagens align laterally with an offset that gives rise to the characteristic D-banding pattern of fibrils. On the other hand, collagen IV, a network-forming collagen, is not post-translationally cleaved. Its end domains serve an important role in network formation. The N-terminal end aligns four molecules and forms a 7S tetramer. On the other end, two molecules form head-to-head assemblies forming an NC1 hexamer. Recent studies show that chloride ions are essential in forming an organized network. Lateral interactions between collagens may also contribute to network assembly (not indicated in this schematic). Schematics are not to scale.

All collagens share a characteristic triple helical structure, formed by a (Gly-X-Y)_n_ repeating sequence in each of the three composite α-chains. However, two key molecular features distinguish collagen IV and fibril-forming collagens that have been shown to impact higher-order assembly. One is the globular C-terminal domain, whose presence inhibits fibril formation and is proteolytically removed from fibril-forming collagens prior to assembly (12). By contrast, the C-terminal non-collagenous domain of collagen IV (NC1) is not removed, and plays a central, chloride-directed role in forming networks (13, 14). The second distinguishing feature of collagen IV is the presence of natural discontinuities (interruptions) in the (Gly-X-Y)_n_ repeating unit within its collagenous domain. The absence or replacement of glycine every third residue prevents continuously triple-helical collagenous domains and promotes local destabilization (15, 16). Interruptions are also present in other non-fibrillar collagens, such as the FACITs, MACITs and MULTIPLEXINs (1). Enhanced flexibility attributed to the presence of interruptions in collagens has been seen using rotary shadowing electron microscopy (17–20), however a detailed study on characterizing the functionality of distinct interruptions is lacking. Because triple-helix interruptions can result from disease-associated mutations in fibril-forming collagens, and because for collagen IV they may play a biological role in providing recognition sites for other macromolecular components in basement membranes (21–23), the ability to interrogate the physical properties of collagen in a sequence-specific manner is needed.

The size of the most abundant mammalian collagen proteins (>300 kDa; >300 nm length) has imposed practical limitations on the structural insight available from conventional approaches. Diffraction-based analysis of molecular structure has been limited to studies of periodic structure within collagen fibrils (24, 25) and to investigations of short triple-helical peptides (~30 amino acids / 10 nm in length) that are amenable to crystallization (23, 26). These and other approaches have provided insight into the molecular determinants of triple helix structure and stability, suggesting for example that imino acids (proline and hydroxyproline) are important for structural stability, and that local interruptions in the (Gly-X-Y)_n_ repeating sequence associated with disease in fibril-forming collagens can disrupt the local structure (16, 23). NMR studies on peptides have also found interruptions to result in increased local chain dynamics (23). Because collagen IV exists as heterotrimers, an additional structural question that arises – beyond whether or not a local sequence possesses a triple-helical structure – is the relative arrangement or stagger among its composite chains (27). Alternative methods for investigating the sequence-dependence of collagen structure in the context of the full-length protein are needed.

Here, we use atomic-force microscopy (AFM) imaging of individual full-length collagen proteins to provide insight into the local determinants of structure. Image analysis provides a map of local flexibility as a function of position along the collagen chain. By applying this technique to collagen IV and to collagen III, we contrast the mechanics of network-forming and fibril-forming collagens at the molecular scale. By mapping the position-dependent flexibility, we relate the local sequence within full-length collagen proteins to the local bending mechanics. Because of the role of chemical environment in regulating collagen supramolecular assembly, we apply this AFM-based flexibility mapping to investigate the effects of distinct chemical environments on flexibility of collagens, globally and in a sequence-specific manner. We interpret these results in light of outstanding physiological questions regarding control of collagen conformations during secretion and its supramolecular assembly in the extracellular space.

## Materials and Methods

### Collagen Sources

Heterotrimeric [α1(IV)]_2_-α2(IV)] collagen IV (113 μg/mL in 0.5 M acetic acid) was purified from Engelbreth-Holm-Swarn (EHS) tumor in lathrytic mouse and was a gift of Albert Ries (28, 29). Rat tail tendon-derived collagen I was purchased from Cultrex (3440-100-01) and is pepsin-treated collagen with a stock concentration of 5 mg/ml in 20 mM acetic acid. Bovine pro-N collagen III (pN-III) was extracted from fetal bovine skin following a previously published protocol (30).

### Sample Preparation for AFM

Collagen was diluted into the desired solution conditions at a final concentration of 0.2 μg/mL, where 50 μL of the diluted sample was deposited and allowed to incubate for 20 seconds on freshly cleaved mica (Highest Grade V1 AFM Mica Discs, 10 mm, Ted Pella). The excess unbound proteins were removed by rinsing with ultra-pure water, and the mica was then dried using filtered air. All proteins were imaged under dry conditions, and the solution conditions of the samples refer to the conditions in which they were deposited onto mica. Previous work suggests that collagen conformations on the mica surface reflect their deposition conditions (10). Imaging was done with an Asylum Research MFP-3D atomic force microscope using AC tapping mode in air. AFM tips with a 160 kHz resonance frequency and 5 N/m force constant (MikroMasch, HQ: NSC14/AL BS) were used.

### Chain Tracing and Analysis

SmarTrace, a custom-built MATLAB code, was used to determine the bending flexibility of collagen chains (10). Persistence length determination by SmarTrace has been extensively validated (10, 31). Chains were traced to subpixel resolution starting from the C-terminus (NC1-chain boundary for collagen IV; non-globular end for collagen III). Homogeneous chain analysis assumes chains to have uniform flexibility (persistence length). The persistence length was found from the dependence of mean squared end-to-end distance, ⟨*R*^2^(Δ*s*)⟩ (Eq. 1), and tangent vector correlation, ⟨cos *θ* (Δ*s*)⟩ (Eq. 2), on the segment length Δ*s*. This model treats the chains as worm-like chains equilibrated in two dimensions, and has previously been shown to describe fibril-forming collagens well in 100 mM KCl, 1 mM HCl (10).

To investigate the sequence-dependence of collagen’s flexibility, the local effective persistence length was determined using Eq. 4 (described in more detail in Supporting Text 1, and Fig. S1). 95% confidence intervals for the estimates of *p*(*s*) were determined using the cumulative probability function of the scaled inverse *χ*^2^ distribution (Fig. S2), as described in Supporting Text 1. The dependence of this estimate on the number of chains is provided in Fig. S3. Analysis of simulated chain images showed that a window of length Δ*s* = 30 nm was the minimum Δ*s* that reliably produced the expected angular distributions for a wide range of persistence lengths tested (Fig. S4). Thus, for a chain of contour length *L*, persistence lengths could be extracted for segments centered from *s* = 15 nm through *s* = *L* − 15 nm.

Simulated images of chains with inhomogeneous flexibility were generated in MATLAB (32) and analysed as described for the experimental images. Chains were generated as described in (10), but with a bending stiffness that varies along the chain and incorporating a “knob” at one end as a directional reference point. Specifically, the chains we simulated here had a total contour length of *L* = 400 nm, interspersing long, stiff regions with short flexible regions (Fig. 3B): *p* = 85 nm (Δ*s* = 79 nm); *p* = 5 nm (Δ*s* = 1 nm); *p* = 85 nm (Δ*s* = 78 nm); *p* = 5 nm (Δ*s* = 2 nm); *p* = 85 nm (Δ*s* = 77 nm); *p* = 5 nm (Δ*s* = 3 nm); *p* = 85 nm (Δ*s* = 76 nm); *p* = 5 nm (Δ*s* = 4 nm); *p* = 85 nm (Δ*s* = 80 nm). The knob was introduced as a uniform-intensity disc of radius 7 nm, centered at the starting point of the simulated chain. Experimentally realistic background noise was included in the images, as described previously (10).

**Fig. 2.**
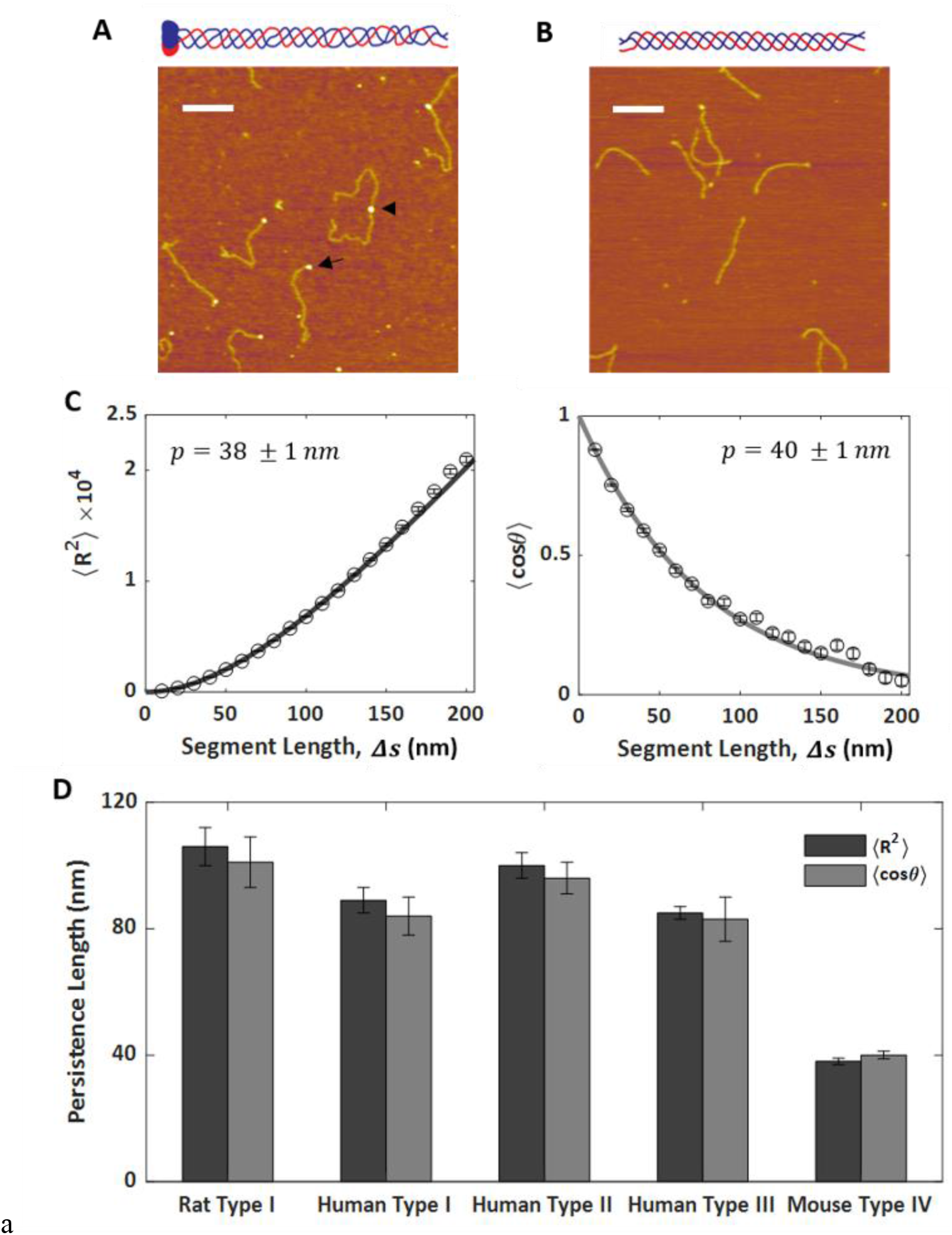
Homogeneous worm-like chain analysis comparing flexibility of collagen IV to fibril-forming collagens. AFM images of **(A)** mouse collagen IV and **(B)** rat collagen I, with schematics of their corresponding molecular structures placed above the AFM images. The black arrowhead and arrow point at an NC1 hexamer and an NC1 trimer of a molecule, respectively. Scale bars = 200 nm. All collagen types were deposited from room temperature onto mica from a solution of 100 mM KCl and 1 mM HCl. **(C)** The persistence lengths were obtained using ⟨*R*^2^(Δ*s*)⟩ (dark gray; equation [1]) and ⟨cos*θ*(Δ*s*)⟩ (light gray; equation [2]) analyses, as shown for the collagen type IV data here. These data are fit well by the predictions of the worm-like chain model. **(D)** Bar plot comparing the persistence lengths of fibril-forming collages to collagen IV. Values of *p* were obtained using the ⟨*R*^2^(Δ*s*)⟩ (dark gray) and ⟨cos 0(Δ*s*)⟩ (light gray) WLC fits. The fibril-forming collagens possess similar persistence lengths of *p* ≈ 90 nm (data reproduced from reference (10)), while collagen IV exhibits a substantially lower effective persistence length of *p* = 39 nm. *N* = 262 collagen IV chains.

**Figure 3.**
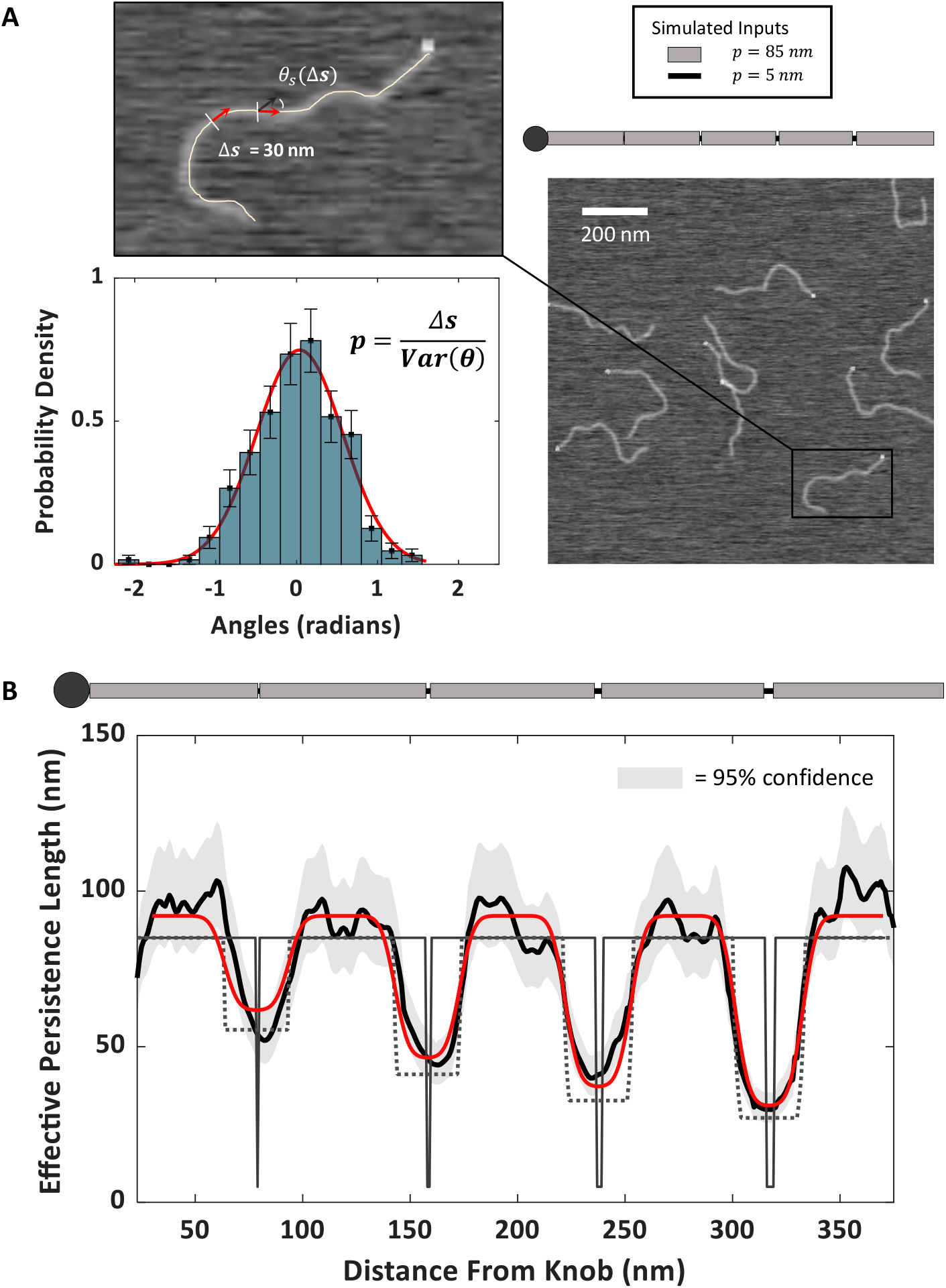
Extracted persistence length profiles from simulated inhomogeneous chains. **(A)** Images of worm-like chains with inhomogeneous flexibility profiles were simulated to validate position-dependent flexibility analysis. For segments of length Δ*s* = 30 nm centered at position *s* along the backbone, the effective persistence length is determined by the variance of angular changes along these segments θ_*s*_(Δ*s*) (equation 4). **(B)** The extracted persistence length profile *p*\*(*s*) obtained by tracing and analyzing *N* = 297 simulated chains (thick black line) shows inhomogeneous flexibility along the simulated chains. The shaded regions represent 95% confidence intervals on the estimates of persistence length. *p*\*(*s*) attains the expected persistence lengths in plateaus longer than Δ*s* = 30 nm and exhibits minima at locations of enhanced flexibility. Minima are broadened with respect to the input *p*(*s*) profile (thin black line), in agreement with the expected effective persistence length profile obtained by convolving a Δ*s* = 30 nm filter with the input *p*(*s*) (dotted line). The two-class model fit (red) recovers distinct persistence lengths of the rigid and flexible regions, returning *p*_r_ = 92 nm and *p*_f_ = 6 nm, respectively (χ^2^_r_ = 1.06). *N* = 297 simulated chains.

### Sequence alignment

The α1 (P02463) and α2 (P08122) amino acid sequences were obtained from UniProt (33). All chain alignments were initiated at the edge of NC1 domain, starting at the first Gly-X-Y unit of collagenous domain of both α chains. Specifics of the alignments are provided in a supporting document. Graphical alignment of the amino acid representation and persistence length profile assumes a length of 0.29 nm / aa in the collagenous domain.

### Variable flexibility model

Each amino acid of an α-chain was assigned a value of “0” if it was within a (Gly-X-Y) sequence or “1” otherwise (within an interruption). Distinct alignments of the three α-chain sequences were tested (supplied as a supporting document), including linear alignment, loop(s) within α2 to better align its (Gly-X-Y) sequence blocks with α1 (including a disulfide-bridged loop (34)), and sequence-similarity alignments (based on a Clustal alignment of the sequences (35), adapted to ensure chain continuity). A flexibility class was assigned at each amino-acid step along the aligned chains, based on whether it was a 0-, 1-, 2- or 3-chain interruption in the triple helix (Table 1). Each of these classes was assigned a local bending stiffness (via a local persistence length) and thus the model assigned a local flexibility to each position along the contour. This model was then fit to the determined effective persistence length profile *p*\*(*s*), in order to determine the persistence lengths *p*_0_, *p*_1_, *p*_2_ and *p*_3_ that best describe each flexibility class. Conversion factors were included in the fitting routine, to convert between primary (aligned) sequence, measured in amino acids, and position along the contour from image analysis, measured in nanometres. Because of variability in identifying the starting position of the chains (found also when tracing simulated chains), a chain-stagger parameter was included, which represented the standard deviation of model chain starting positions. Finally, because we do not know where the collagenous domains start (for example, their starting position may be obscured by the NC1 domain), an offset parameter was included to linearly shift the modeled chains with respect to the traced chains. The fitting procedure is described in detail in the Supporting Text 2.

**Table 1:** Physical classes of interruption.

| Flexibility class | $\alpha 1$ interruption? | $\alpha 2$ interruption? |
| --- | --- | --- |
| Triple helix ($p_0$) | 0 | 0 |
| One-chain interruption ($p_1$) | 0 | 1 |
| Two-chain interruption ($p_2$) | 1 | 0 |
| Three-chain interruption ($p_3$) | 1 | 1 |

Modelling the simulated chains (Fig. 3) required a simpler physical model containing only two classes: rigid *p*_r_ and flexible *p*_f_, located along the chains as defined by the inputs to the simulations. No length conversion factors were necessary because the simulated chain profile positions were defined in nanometres. Chain-stagger and offset parameters were necessary to align the model results with the traced and analysed simulated-chain*p*\*(*s*) profile.

## Results and Discussion

Most studies of collagen mechanics have focused on fibril-forming collagens such as collagen I, which associate laterally to form higher-order structures as shown in Fig. 1 (8). Considerably less is known about the mechanics of collagen IV, which forms the scaffold of basement membranes. We first quantified flexibility differences between these types of collagens by analyzing images of many individual collagen molecules.

### Atomic Force Microscopy Images of Collagen

We imaged mouse heterotrimeric [α1(IV)]_2_-α2(IV)] collagen IV molecules (also referred to as protomers), derived from the matrix produced from Engelbreth-Holm-Swarn (EHS) tumor, using atomic force microscopy (AFM). Because it does not require a metal replicate but instead directly images molecules deposited on a surface, AFM is free from potential artefacts of rotary shadowing electron microscopy (EM), which has previously been used for imaging conformations and binding interactions of collagen IV (4, 17, 18, 36–42). With appropriate imaging conditions, AFM offers superior spatial resolution than the platinum nanocrystallite size of 1-2 nm in EM (17), and our studies avoid the use of glycerol, which has been shown to affect collagen IV assembly (36). Although imaging in liquid conditions is possible with AFM, here, we deposited the collagen from the specified solution conditions and dried prior to imaging. This follows previous work that used AFM to image and quantify the flexibility of collagen types I, II and III (10, 31, 43), to observe interactions between the NC1 domains of collagen IV (44, 45), and to observe binding sites of laminin on collagen IV (46). For the conditions used in this study (>100 mM ionic strength deposition prior to drying), numerous statistical measures demonstrate the collagen to be equilibrated on the mica surface (10).

An example AFM image of collagen IV deposited from acidic solution (100 mM KCl, 1 mM HCl) is shown in Fig. 2A, along with an image of collagen I for comparison. The acidic conditions preclude lateral assembly of both fibril-forming and collagen IV molecules (10, 36), while the presence of chloride ions can induce oligomerization of collagen IV protomers towards network formation (14, 44). Both proteins possess long chains, representing the collagenous domains. The contour length for EHS-derived collagen IV protomers is 360 ± 20 nm (*N* = 262), in agreement with previous reports (36) and, as expected, longer than the ~300 nm contour length of fibril-forming collagens (17). Similar to previous images recorded using rotary shadowing EM (17, 36), we observe by AFM that the collagen IV protomer is capped by a globular domain, known as the C-terminal NC1 domain. Evident in the AFM image are two kinds of molecules: single protomers and dimers of protomers linked end-on by NC1 hexamers.

Visual comparison of collagen I and collagen IV images suggests that the triple helix of collagen IV is more flexible: collagen I bends on a long length scale, while collagen IV exhibits more frequent and shorter-range bending fluctuations.

### Conformational analysis of collagen IV as a homogenous polymer

To quantify the flexibility of the collagenous domain of collagen IV, we applied our previously developed chain tracing and analysis algorithm, SmarTrace (10). SmarTrace traces and provides conformational analysis of imaged chains, allowing for the implementation of polymer physics tools to determine their mechanical properties. To analyze collagen as a homogeneous polymer, we traced chains collected from AFM images and randomly segmented the contours into non-overlapping pieces of different segment lengths. We calculated the mean squared end-to-end distance ⟨*R*^2^(Δ*s*)⟩ and the mean correlation of the beginning and ending unit tangent vectors (given by their dot product) 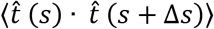 for all segments of length Δ*s* within the pool of collected chains. These were fit with the predictions of the inextensible worm-like chain (WLC) model to estimate persistence length (10, 47):

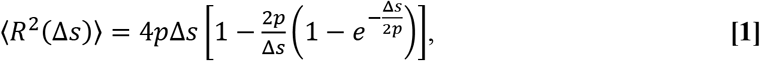

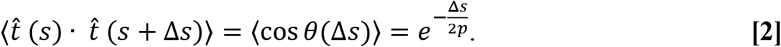

These equations assume the polymers to be equilibrated in two dimensions, which has been shown to be the case for collagens deposited under the solution conditions used here (10). While we cannot test this explicitly, we assume that the persistence lengths determined from these two-dimensional images represent the bending flexibility of collagen in three dimensions, an assumption that holds reasonable agreement with previous collagen flexibility measurements (see discussion in (10)) and that has been clearly demonstrated for other systems like DNA (47, 48). The collagen IV results for both ⟨(*R*^2^(Δ*s*)⟩ and ⟨cos *θ*(Δ*s*)⟩ appear to be well described by the WLC model (Fig. 2C), from which we find a persistence length of *p* = 39 ± 2 nm. To our knowledge, this is the first report of a net persistence length of collagen IV.

The persistence length of collagen IV is significantly less than that of fibril-forming collagens deposited under the same solution conditions (Fig. 2D). Our earlier work revealed that the persistence lengths of different fibril-forming collagens (I, II and III) are very similar, all falling within ~10% of *p* = 90 nm under these solution conditions and being well described as semi-flexible polymers (10). By contrast, the persistence length of collagen IV is less than half of this value, reflecting a more flexible protein. This stark difference in bending flexibility between collagen IV and fibril-forming collagens contrasts with their very similar thermal stabilities (49). (This is achieved in part via more extensive proline hydroxylation in collagen IV, which compensates for the presence of interruptions (49).)

This striking difference in flexibility likely relates to the interruptions that characterize collagen IV molecules. The triple helix of collagens is defined by a repetitive (Gly-X-Y)_n_ sequence, in which the Gly is obligatory to form a stable triple-helical structure. The collagenous domain of collagen I is 96% triple helical (with non-helical regions confined to its telopeptide ends (50)) while in collagen IV it is approximately 80% triple helical, with interruptions to the (Gly-X-Y)_n_ sequence occurring throughout the chain (Fig. S5). This distinction is shown schematically above the AFM images in Fig. 2. Because unstructured polypeptide chains are extremely flexible with *p* < 1 nm (51), we expect interruptions in the triple-helix-defining sequence to significantly enhance flexibility. This expectation is consistent with our finding of a lower persistence length of collagen IV and with the visual comparison of images of collagen IV and collagen I chains (Fig. 2A, B).

From the example chain images in Fig. 2A, it appears that the flexibility of the collagen IV molecule varies along its contour and is more flexible away from the NC1 domain, *i.e*., towards the N terminus. In a study using rotary shadowing electron microscopy to examine collagen IV, Hofmann *et al*. found that it exhibits regions of variable flexibility along its length (17). Thus, although the homogeneous worm-like chain model provides a good fit to our experimental data (Fig. 2C), we sought to assess the heterogeneity of flexibility along the collagen IV triple helix.

### Position-Dependent Flexibility Analysis

To address structural variability, and to evaluate the contributions of interrupted regions to the overall flexibility of collagen IV, we extended the SmarTrace analysis to include determination of the sequence-dependent chain flexibility. To do so, we use the variance of tangent angles around each position s along the chain backbone to quantify the bending stiffness at that location. This approach was developed by Hofmann *et al*. to analyse the position-dependent flexibility of collagens (17). This method complements other AFM image analysis methods developed to quantify discontinuous mechanical properties of DNA (52) or to map sequence-dependent properties of individual proteoglycans (53), and is expected to be more appropriate for flexible molecules than a position-dependent stiffness analysis developed for more rigid biological filaments (54).

The flexibility of a filament at a position *s* along its contour is described by its local bending rigidity *α*(*s*). The bending rigidity is related to the persistence length via the thermal energy, *k*_B_*T*:

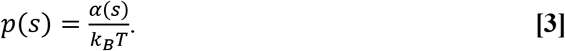

The persistence length thus describes the flexibility of a chain with a given bending stiffness, at a given temperature. Practically, we are limited to examining the flexibility of a chain over a segment of finite length Δ*s*, which we define centered at position *s*; we use the term “effective persistence length” to denote this averaged nature of the response, *p*\*(*s*;Δ*s*). The effective persistence length of a segment of length Δ*s* is determined from the variance of the tangent angles *θ*s between the ends of the segment centered at *s*:

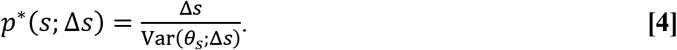

The analysis methodology was validated through tests on simulated chains. We generated AFM images of chains with inhomogeneous flexibility profiles, traced the chains using SmarTrace, and analysed the resulting contours using Eq. 4. The chains were simulated as previously described (10), here, incorporating a position-dependent bending stiffness and a “knob” at one end (which served as a directional reference point, analogous to the NC1 domain of collagen IV). Fig. 3A shows an example of a simulated image, along with a schematic of the inhomogeneous flexibility profile imposed in the simulations (long regions of relatively stiff chains with *p*=85 nm, interspersed with very short regions with *p*=5 nm). The short regions of substantially increased flexibility are not visually apparent in the simulated images, but are clearly seen in the effective persistence length profile determined from the traced chains (Fig. 3B). A notable difference between the input and extracted persistence length profiles is the apparent broadening of the flexible regions: *p*\*(*s*;Δ*s*) exhibits much broader wells than the input *p*(*s*) profile. This is the expected result of the analysis: the Δ*s*=30 nm segment length is substantially longer than the regions of flexibility in our simulated chains and acts to some degree as a low-pass filter of flexibility along the chain (though we stress that *p*\*(*s*) is *not* simply the mean of *p*(*s*) over a 30-nm window; see Supporting Text 1 and Fig. S1). Fig. 3B includes the expected effective persistence length profile, which takes into account this convolution between the ideal chain and the 30-nm filter. This expected profile agrees well with *p*\*(*s*;Δ*s*) obtained from analyzing the simulated images. While a shorter segment length should provide more localized information on sequence-dependent flexibility, our validation tests found Δ*s*=30 nm to be the minimum length that returned reliable estimates of *p*\*(*s*) from simulated chain images (Fig. S4). Thus, Δ*s*=30 nm was used for the analysis presented herein.

Having validated the persistence-length mapping algorithm, we applied it to experimental images of collagen IV. The effective persistence length profile of collagen IV is shown in Fig. 4. From this result, it is obvious that the flexibility of collagen IV varies markedly along its length, with the N-terminal half of the molecule being significantly more flexible than the region closer to the NC1 domain. Similar flexibility trends are observed whether collagen chains are traced starting at the NC1 domain (Fig. 4) or in the reverse direction from the 7S domain (Fig. S6). Our flexibility profile agrees well with that of Hofmann *et al*., who also observed increased flexibility towards the N-terminus of collagen IV (17).

**Figure 4.**
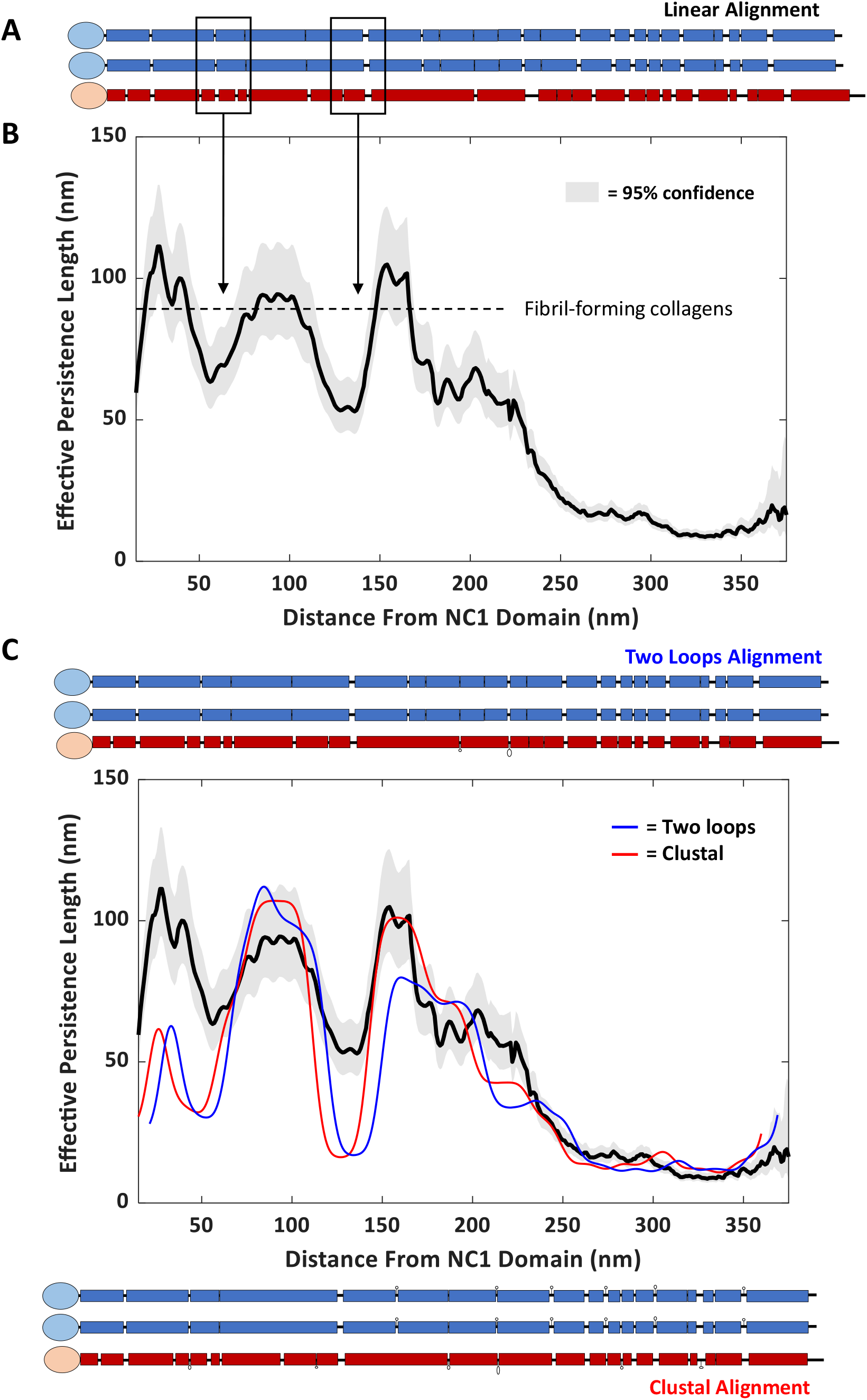
Position-dependent flexibility profile of collagen IV. **(A)** Schematic representation of the α1 and α2 amino acid sequences from mouse collagen type IV. Rectangles indicate regions of the sequence containing triple-helix-competent sequences with (Gly-X-Y)_n_G, *n* ≥ 4. Interruptions of this repetitive sequence are indicated by thinner lines, and occur more frequently towards the N-terminus of the chains (right side of the schematic). **(B)** Position-dependent persistence length map of collagen type IV deposited from 100 mM KCl, 1 mM HCl. The dashed line indicates the persistence length of continuously triple-helical fibril-forming collagen molecules (10). Shaded curves represent 95% confidence intervals on the effective persistence length estimate *p*(*s*; Δ*s*). The profile was calculated from *N* = 262 chains. **(C)** Four-class flexibility model using two loops (blue) and Clustal (red) alignments. Alignments differ in the amount of amino acids that loop out and do not participate in the main backbone, as shown schematically above and below the plot. The persistence length profile is aligned with the amino acid sequence representations using model outputs for best-fit offset (nm) and nm/aa conversion. The two loops (Clustal) alignment is best fit with persistence lengths of *p*_0_ = 105 (109) nm; *p*_1_ = 83 (86) nm; *p*_2_ = 21 (42) nm; *p*_3_ = 1.9 (2.0) nm, with χ^2^_r_ = 12.6 (12.8). Best-fit persistence lengths of these different structural classes – particularly *p*_1_ and *p*_2_ – differ for other chain alignments (Table S1).

To gain sequence-dependent insight into the flexibility profile requires aligning this map with the collagen IV sequence, which was unavailable to Hofmann *et al*. (17). An amino acid representation of the full sequence of mouse [α1(IV)]_2_α2(IV) collagen IV is shown in Fig. 4A. The α-chain polypeptide sequences have been aligned from the beginning of the collagenous domain beside the NC1 domain (from the first Gly-X-Y). The blue and red boxes correspond to segments of α1(IV) and α2(IV), respectively, that have the (Gly-X-Y)_n_ sequence required to form a triple helix. In this representation, we have assumed a minimum length of (Gly-X-Y)_4_G to be sufficient to form a triple helix. This assumption of *n*=4 does not strongly affect the representation (Fig. S7). The thinner lines in Fig. 4A represent interruptions in the triple-helix-competent (Gly-X-Y)_n_ repetitive sequence.

The position-dependent flexibility profile of collagen IV aligned with the amino acid sequences is shown in Fig. 4A-B. An examination of the sequences of the α-chains reveals that most of the interruptions lie in the N-terminal half of the protein, where we also see an increase in flexibility (lower persistence length). Maxima in local persistence length align well with extended triple-helical stretches in all three chains, found approximately 30, 90, and 160 nm from the NC1 domain. Within these plateau regions, collagen IV’s effective persistence length is *p*\* ≈ 95 nm, the persistence length found for continuously triple-helical collagens (see Fig. 2) (10). Between these maxima, minima in persistence length are found approximately 60 and 130 nm from the NC1 domain. These two locations align well with regions of the sequence in which both α1 and α2 chains possess interruptions in the (Gly-X-Y)_n_ sequence. This suggests that interruptions of the triple-helix-forming sequence in all three α-chains of α1(IV)2-α2(IV) strongly impact the flexibility of the molecule, apparently more so than having interruptions in only one or two of the constituent chains.

### Model for interpreting position-dependent flexibility

To gain a deeper structural interpretation of the persistence length profile, we implemented a simple physical model to describe the sequence-based flexibility of a collagen containing interruptions (Supporting Text 2). This model assumes that the local flexibility of collagen can be completely described by how many of the three component chains contain a triple-helix-compatible sequence at that location. To implement the model, each amino acid in each of the α1 and α2 chains was assigned a value of either “0” (contained within triple-helix-compatible sequence (Gly-X-Y)_n_) or “1” (within an interruption). The three α-chain sequences were aligned starting from the edge of the NC1 domain, and different chain alignments were tested. These include a linear, sequence-based alignment (Fig. 4A and Fig. S8); nonlinear alignments that include previously proposed intrachain loops within α2 (34, 55, 56) and improve the registration of triplehelix-forming sequences; and alignments arising from a Clustal-based analysis of sequence similarity between α1 and α2 chains (35), which require incorporation of copious loops and bulges to confer backbone continuity.

For each chain alignment, a flexibility class was assigned at each amino-acid step along the backbone based on whether it was a 0-, 1-, 2- or 3-chain interruption in the triple helix (Table 1). Each of these classes was assigned a local bending stiffness (via a local persistence length) and thus the model assigned a local flexibility to each position along the contour. This model was then fit to the determined persistence length profile *p\**(*s*;Δ*s*), in order to determine the persistence length *p*_0_, *p*_1_, *p*_2_ and *p*_3_ that best describes each flexibility class. The approach was validated by application to the simulated chains: here, a two-class model (rigid or flexible) recovered the distinct persistence lengths of the rigid and flexible regions and described the data well (*p*_r_ = 92 nm; *p*_f_ = 6 nm; χ^2^_r_ = 1.06). A slight overestimate of persistence lengths is expected for finite samples (Supporting Text 1). Agreement between the four-class flexibility model and the measured persistence length profile *p\**(*s*;Δ*s*) of collagen IV would indicate that this physical classification is a valid approach to describing the variable flexibility of collagen IV. By contrast, disagreement would indicate that other factors (such as the influence of X and Y identity on flexibility, length of interruption, and location with respect to interruptions (extended triple-helix tracts) may be of key importance for quantifying the structural properties of this protein.

This model of a physically heterogeneous collagen IV captured positional variations in flexibility along the chain, comparing favorably to the measured *p\**(*s*) profile (Figure 4C). The model returned a value for triple helix local persistence length of *p*_0_~100-160 nm for all distinct α-chain alignments tested (Table S1). This estimate of *p*_0_ is somewhat larger than the net persistence length of continuously triple-helical collagens, found by treating the chains as homogeneous (Fig. 2D), but is consistent with local values of *p*\* determined along their backbones (see below). Also consistent among chain alignments was the finding that *p*_2_ < *p*_1_. In other words, having an interruption in two of the chains leads to a more flexible structure than having a one-chain interruption.

Some of the chain alignments resulted in fits of the variable flexibility model that were not minimized within the specified parameter range: for example, a linear chain alignment (aligned based only on primary sequence) produced for an overlapping interaction *p*_3_ = 200 nm, the maximum value allowed for fitting, a value that represents an unphysically rigid structure of three polypeptide chains that lack any triple-helix-forming sequence. Alignments that returned this high value, however, were the only ones that captured the rigidity in the collagenous region adjacent to the NC1 domain (e.g. Fig. S8).

Alignments that optimized the fit within their parameter bounds returned an optimal length scaling of 0.29 nm / amino acid, commensurate with triple helix lengths from collagen diffraction analysis (23, 57). These alignments also returned the smallest value for *p*_3_ ≈ 2 nm, *i.e*., that an overlapping interruption is the most locally flexible structure within the collagenous domain of collagen IV. These optimal alignments require at least some outward loops in the longer α2 chain (e.g. “Two loops” alignments) and perhaps extensive minor loops and bulges along the length of each of the three α-chains (“Clustal” alignments).

In some regions, the variable flexibility model underestimates the rigidity, all located in the C-terminal half of the protein (Fig. 4C). The region adjacent to the NC1 domain exhibits a rigidity similar to the other extended triple-helical regions, yet the model significantly underestimates the persistence length, both of the plateau at *s* ≈ 30 nm and of the adjacent minimum at *s* ≈ 50 nm. These model results imply that the overlapping interruption closest to the NC1 domain possesses a distinctly enhanced stability not captured by our simple physical model. Enhanced stability in this region could be conferred by posttranslational glycosylation of hydroxylysines within this overlapping interruption (58–60). Unfortunately, a glycosylation map of collagen IV lacked sequence coverage in this region (61), so this remains a speculative, yet testable, prediction.

The alignment of the three chains of the α1α1α2 mouse collagen IV collagenous domain is unknown. A disulfide-bridged loop within α2 has been proposed, which helps better to align triple-helix-competent sequences in the chains (34, 55); this is included as the larger, N-terminal loop in our “two-loops” alignment (Fig. 4 and Supporting Information). Most of our chain alignments also agree with the determined alignment of α chains at an integrin-binding site in human collagen IV (Table S1), the only location for which chain alignment in collagen IV has been determined (27). To validate or further constrain chain alignments will require improvements in spatial resolution from this AFM imaging and chain tracing approach and/or complementary approaches such as proteolysis to identify unstably structured regions within the collagenous domain (62).

In our alignments, we ignored the register of the three chains, since the register has been identified at only one location and only in human collagen IV (27). We tested a staggered alignment of the three chains that included this identified local alignment, but it failed to produce an optimized fit (Table S1). It is highly likely that some of the interruptions alter the local register of the three chains (such that a leading chain at some locations may become the middle or trailing chain at others) (16), and that one “universal” registration does not apply along the entire collagenous domain.

It is also possible that the variable flexibility model is insufficient to completely describe the measured flexibility profile because of its many simplifying assumptions. Beyond stabilization by posttranslational modifications and local variations in chain stagger, there are other attributes that may be of direct relevance to local stability and flexibility. For example, we have ignored the length-dependence of interruptions when defining flexibility class. Short one-chain interruptions may act as “stammers”, serving to alter the local register among chains instead of breaking the triple helix structure (16). Single glycine substitutions may introduce local kinks in the structure (63). Longer interruptions may destabilize flanking triple-helical regions, leading to increased breathing of the adjoining triple helix and a lower persistence length (64). The local sequence within and flanking the interruption also may alter its stability (23). And finally, this physical model ignores all sequence-dependent attributes of not only the interruptions but also of the triple helix itself, by classifying each amino acid purely by whether or not it is found within a (Gly-X-Y) sequence. We next sought to test the assumption of mechanical homogeneity within the triple helix, by mapping the flexibility profile of a fibril-forming collagen.

### Triple helix mechanical inhomogeneity: collagen III

To probe the variability of bending stiffness along the triple helix, we imaged the pN-III molecular variant of bovine collagen III using AFM (Fig 5A). Collagen III is a homotrimeric [α1(III)]_3_] fibril-forming collagen devoid of interruptions in its ~300-nm-long triple helical domain. The pN-III variant retains its N-terminal propeptide (65), providing us with a directional reference point for position-dependent flexibility analysis.

**Figure 5.**
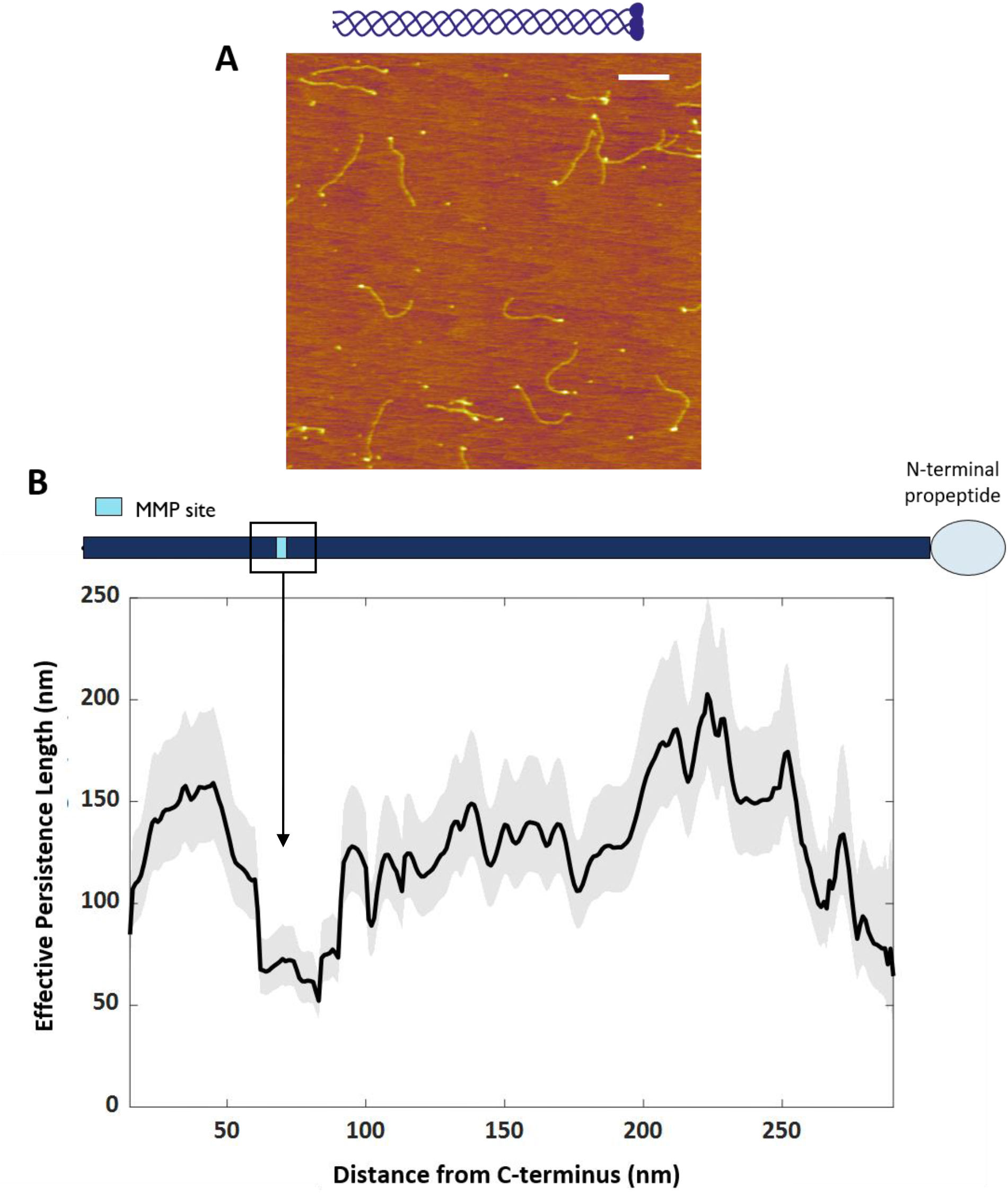
Position-dependent flexibility profile of collagen III. **(A)** AFM image of bovine collagen pN-III deposited from room temperature onto mica from a solution of 100 mM KCl and 1 mM HCl. Scale bar = 200 nm. **(B)** Position-dependent persistence length map of collagen III aligned with its amino acid sequence representation. This collagen is continuously triple-helical and therefore is represented as a single bar with the MMP site marked. Shaded curves represent 95% confidence intervals on the effective persistence length. The profile was calculated from *N* = 267 chains.

We found that the flexibility of a triple helix varies significantly along its contour, although less than along the interruption-containing collagen IV (Fig. 5B). This provides direct evidence for distinct mechanical properties of different sequences within the collagen triple helix. Our results show considerably more variation in local flexibility along the backbone of collagen III than previous rotary-shadowing EM measurements on this molecule (17), perhaps due to the aforementioned higher resolution provided by AFM and/or the choice of segment length used to determine the effective persistence length. Our flexibility map shows that the effective persistence length ranges from approximately 60-180 nm in different regions of collagen III. Notably, the larger values are similar to values returned for *p*_0_ of the triple helix in our model fitting, above, which ranged from 100-160 nm, and are longer than the net persistence length of continuously triple-helical collagens (Fig. 2D). In the context of our physical model, this has additional implications. The range of *p*\* found for collagen III suggests that the assumption of a single persistence length *p*_0_ to characterize all triple-helical regions of collagen IV is an oversimplification. However, the model-extracted values of *p*_2_ and *p*_3_ are significantly shorter than the lowest values of *p*\* along the collagen III chain, with values of *p*_1_ similar to or less than the most flexible regions of collagen III. It is possible that two chains possessing a (Gly-X-Y)_n_ sequence may be able to induce a triple-helix-compatible structure in the third chain, leading to a value of *p*_1_ similar to regions of the collagen III triple helix. Overall, we can conclude that the enhanced flexibility of collagen IV is dominated by the effects of interruptions, while the variable flexibility along collagen III implies that sequence does influence mechanics at the molecular level.

Regions of greatest flexibility along pN-III collagen are found near the N-terminus (which contains conformationally distinct regions (66, 67)) and, intriguingly, at the matrix-metalloprotease (MMP) site (Fig. 5B). This site is the target of proteolysis by MMPs and is of key importance for extracellular matrix remodelling, e.g. during embryonic development and cancer metastasis. To our knowledge, ours is the first demonstration of enhanced *mechanical* flexibility at the MMP site. Our ability to detect this feature, while previous studies using EM did not (17), may arise from the higher-resolution imaging of AFM, and from our directional tracing from the C-terminus towards the N-terminus. When we traced the chains starting at the N-terminus, the contrast in flexibility around the MMP site was diminished (Fig. S9), perhaps resulting from challenges in identifying the start of the triple helical domain in the presence of the N-propeptide (66).

The MMP site has previously been identified as possessing distinct characteristics within the collagenous domain. Intriguingly, the sequence in this MMP region of collagen I has been found – in peptides – to exhibit structural tautomerism, transitioning between a triple helix and β1-bend structure (68). It is interesting to speculate that this structural flexibility may underlie the enhanced bending flexibility revealed here through AFM imaging. The MMP region also exhibits enhanced sensitivity to proteolysis by trypsin and has been suggested to have an enhanced propensity to unwind (69–75). We propose that this structural instability is what gives rise to the enhanced bending flexibility in this region: three weakly interacting chains are predicted to bend more easily than a tightly wrapped triple helix. This would produce a lower local persistence length.

The decreased triple-helix stability in the MMP region of fibril-forming collagens has been attributed to a low imino-acid content (72, 76, 77). Thus, we examined how local imino-acid content correlates with flexibility (Fig. S10, S11). We saw no obvious correlations between imino acid content and flexibility along the collagen III contour (Fig. S10A), which was confirmed by a linear correlation coefficient of *R*^2^ = 0.017. Although an anticorrelation between imino acid content and flexibility in the C-terminal half of collagen IV might be inferred by visual comparison of these profiles (Fig. S11B), quantitative analysis revealed no statistical correlation between these quantities, neither for the full length of the chain (*R*^2^ = 0.084) nor for the first 200 nm from the NC1 domain (*R*^2^ = 0.15). Thus, a simple picture of imino acids enhancing rigidity (17, 78) does not apply here. Incidentally, recent work has suggested that imino acids may, instead, enhance flexibility of triple helical regions (79). While the anticorrelation we observe in some regions supports this conclusion, we do not find this to be the case globally along the collagen backbone. Thus, while imino acids may contribute to local flexibility, at present our data suggests that they are not the main driver of bending flexibility of the triple helix.

We also looked for correlations between the locations of the imino acid within the (Gly-X-Y) triplet and the flexibility. This is because most prolines in the Y position are 4-hydroxylated (while most in the X position are not, with a small fraction 3-hydroxylated) (61, 80–82). Hydroxyproline increases the thermal stability of the triple helix (83, 84), and we thus wished to determine if its presence correlated with increased triple-helix bending rigidity. As seen in Fig. S10B, we find no correlation between flexibility and Y-positioned prolines. X-positioned prolines can enhance conformational flexibility (within Gly-Pro-Hyp triplets), and have recently been correlated with enhanced thermal stability of collagens (79). However, we also find no correlation between local flexibility within collagen III and the local density of X-positioned prolines (Fig. S10B).

The physical model we have developed to assess the structural relationships between interruptions and flexibility could be adapted to study sequence-dependent influences on flexibility in collagens lacking interruptions, such as collagen III. At present, our flexibility analysis is limited by accuracy in chain registry and has a ~100-amino-acid resolution (30 nm filter window) (Supporting Text 2), but improvements in imaging and tracing algorithms should be able to push this to finer length scales. Even with the current resolution, this approach should allow initial comparisons of bending flexibility with measured and predicted variations in local helical symmetry, thermal stability, sequence content and post-translational modifications. Such comparisons would provide important links among composition, structure and mechanics of collagen, and would offer a further means to connect high-resolution studies on triple-helical peptide constructs with their *in-situ* properties in the full-length proteins (23, 85). Such knowledge of the breadth of properties exhibited by continuously triple-helical structures would enable elucidation of the effects of glycine substitutions and of sequence interruptions on the molecular structure and mechanics of collagen (63, 86, 87).

### Physiological implications of collagen IV flexibility

Thus far, our analysis has determined collagen IV properties only when deposited from acidic solution conditions. These conditions (pH ~ 3) are more acidic than the varying physiological conditions during biosynthesis and secretion. Previous studies of collagen IV flexibility also deposited the protein from very non-physiological conditions (25 mM acetic acid, 50% glycerol) (17). Thus, we sought to investigate its flexibility in more physiologically relevant conditions.

Intracellularly, folded proteins are transported in secretory vesicles with low pH and chloride concentrations relative to the cellular exterior (88, 89). Upon secretion, collagen’s chemical environment changes to one with a higher chloride concentration and neutral pH. This environmental switch is important for triggering assembly into higher-order structures: NC1 interactions strengthen at neutral pH and higher chloride concentrations (42, 90, 91), supporting collagen IV assembly into networks, and neutral pH and chloride favor lateral assembly of fibril-forming collagens (92, 93). Environmental conditions such as pH and ionic strength have also been found to influence the conformations of fibril-forming collagen molecules (10, 43). These observations raise the question of how much the conformations of the collagenous domain of collagen IV are controlled by chloride and pH. We posited that the pH and chloride gradient experienced during secretion might affect the flexibility of collagen IV, conferring a shorter persistence length / enhanced flexibility early in the secretory pathway, which would impart a lower free energy cost for compact configurations.

To test this hypothesis, we analyzed collagen IV under two additional solution conditions of pH and Cl concentrations. To represent vesicular solution conditions, we took an extreme limit by eliminating the chloride ions and used a sodium acetate buffer at a slightly acidic pH [150 mM NaOAc, 25 mM Tris-OAc, pH 5.5]. TBS [150 mM NaCl, 25 mM Tris-Cl, pH 7.4] served as an extracellular proxy because of its relatively high chloride content and neutral pH.

AFM images of collagen IV molecules deposited from each of these two solution conditions are shown in Figs. 6A-B. Visually, the collagen conformations look similar in both images, and appear similar to those found in the previous condition of 100 mM KCl, 1 mM HCl (pH ~3) (Fig. 2A). It is important to note that, although collagen was deposited from a chloride-free environment in the AFM image in Fig. 6B, there are still some head-to-head assemblies. This is most likely due to the presence of sulfilimine cross-links in the NC1 hexamer, post-translational modifications that endow mechanical strength and stability to the collagen IV network (94). The presence of such cross-links protects the dimers from dissociation in a chloride-free environment. We do not observe any evidence of higher-order association or network formation in our images in these conditions, likely due to the low solution concentration of collagen IV used for deposition (0.2 μg/mL) (36).

**Figure 6.**
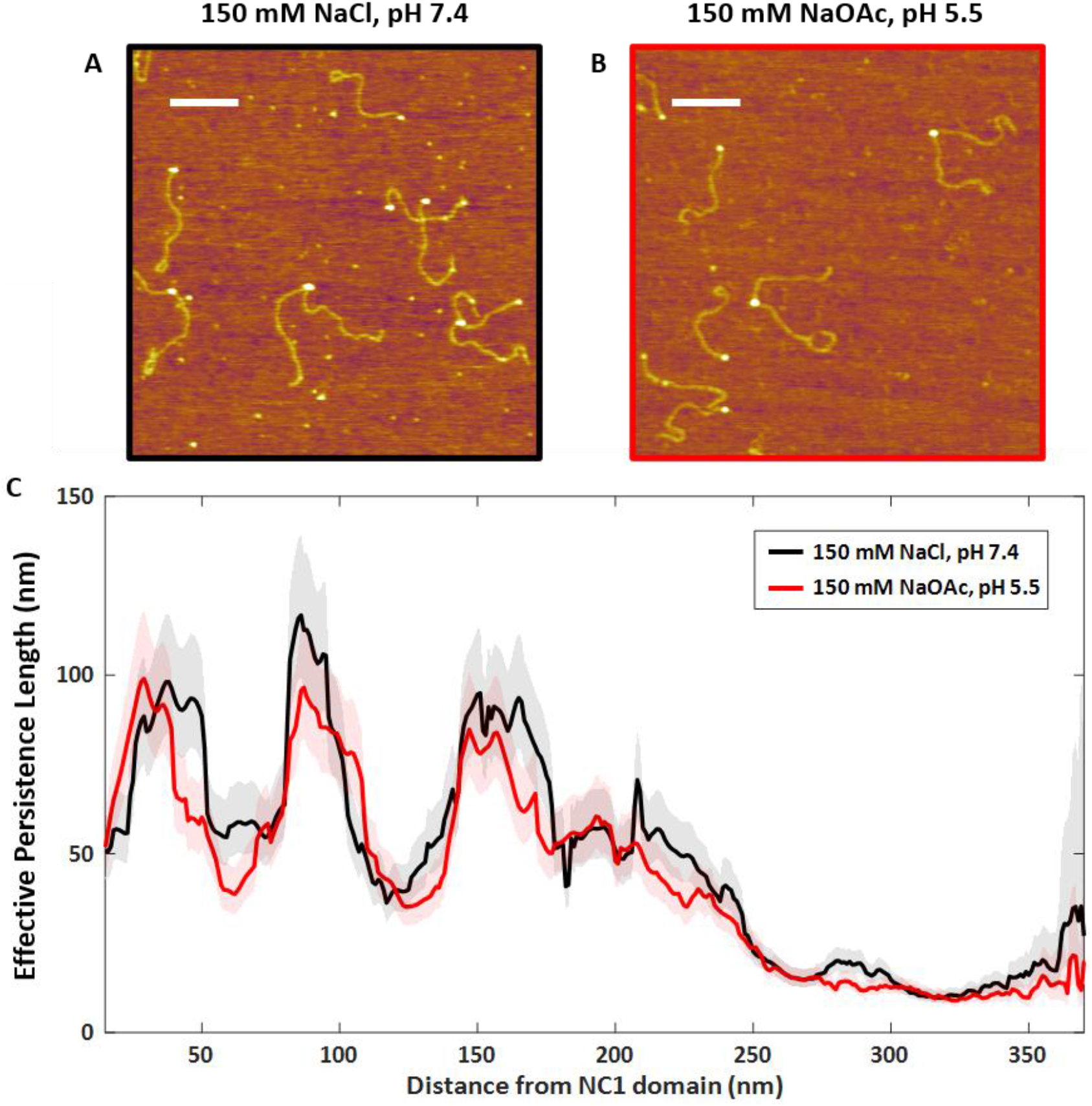
Effect of chloride and pH on the flexibility of collagen IV. Inspired by the changes in the chemical environment along collagen’s secretion pathway, we imaged collagen IV under two solution conditions, **(A)** TBS [150 mM NaCl, 25 mM Tris-Cl, pH 7.4], and **(B)** sodium acetate buffer at a slightly acidic pH [150 mM NaOAc, 25 mM Tris-OAc, pH 5.5]. Scale bars = 200 nm. **(C)** Position-dependent flexibility map of collagen type IV. The shaded curves represent 95% confidence intervals on the effective persistence length estimates *p*(*s*; Δ*s*). The profiles were calculated from *N* = 272 (TBS) and *N* = 273 (NaOAc) chains.

Surprisingly, we find similar overall flexibility among both buffered solution conditions (net persistence lengths *p* = 43 ± 3 nm for TBS and *p* = 41 ± 3 nm for NaOAc; Fig. S12) and the previous acidic solution (p = 39 ± 2 nm for 100 mM KCl + 1 mM HCl). Collagen IV exhibited no global curvature in any of these conditions (data not shown). Therefore, collagen IV flexibility is not tuned by changes in pH and chloride concentrations that mimic those experienced during secretion.

The net persistence length of a collagen molecule can be used to estimate its average size in solution, which we can compare with the size of vesicles that have been proposed to transport collagens from the endoplasmic reticulum (ER) to the Golgi. To do so, we assume that collagen molecules adopt a random coil conformation in solution whose extent is approximated as a sphere with a radius given by the radius of gyration, *R*_g_. We find that collagen IV is predicted to be considerably more compact 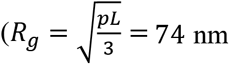, assuming a contour length *L*=410 nm and *p*=40 nm) than the interstitial, fibril-forming collagens (*R_g_* = 95 nm, with *L*=300 nm and *p*=90 nm). Thus, in spite of its longer length, collagen IV would pose less of a size burden to the secretory machinery of the cell if it were to be transported in small vesicles. The 150-190 nm diameter of such random coils is significantly less than the 300-nm length dimension commonly stated as required for transport, determined by (incorrectly) assuming fibril-forming collagens to act as rigid rods. Nonetheless, both collagen IV and fibril-forming collagens would require additional compaction to fit in the standard COPII vesicles involved in ER-to-Golgi transport (60-90 nm diameter) (88, 95).

While it is generally accepted that collagens are *not* trafficked in such small vesicles, there remains considerable uncertainty about the mechanism used for procollagen trafficking during secretion. To our knowledge, there is no evidence of any collagen-associated proteins such as Hsp47 acting to compact collagen to facilitate loading into ER exit vesicles. In fact, some studies have found collagens IV and VII to be transported from the ER to Golgi in larger vesicles with 400 nm diameter (96, 97), into which fibril-forming and collagen IV could easily be accommodated. However, recent work has found that direct trafficking of procollagen I from the ER to Golgi occurs and that these large vesicular carriers are not required (98). In this case, collagen (in)flexibility would not pose a strong energetic cost on its transport to the Golgi, consistent with our finding that collagen flexibility is not tuned by chemical environmental changes experienced during secretion.

As shown above, collagen IV should not be viewed as a uniformly flexible structure, so again, we performed a position-dependent analysis. We sought to determine how both triple-helical and interrupted regions of the collagenous domain of collagen IV are affected by chloride and pH. These regions have been implicated in assembly into higher-order networks (40), and we posited that their stability and flexibility would vary between intracellular (lower Cl^-^ and pH; assembly-disfavouring) and extracellular (higher Cl^-^ and pH; assembly-favouring) conditions. Specifically, because the triple helix is thought to loosen and destabilize at lower pH and Cl^-^ (albeit at significantly lower ionic strengths than used here) (10, 99), we anticipated that a similar destabilization would enhance microunfolding of the triple-helical regions of collagen IV. This would be seen by a shortening of the high persistence length plateaus in sodium acetate relative to TBS, and a concomitant broadening of the minima.

Surprisingly, we find that the flexibility profile of collagen IV is affected very little by these changes in pH and chloride. Fig. 6C shows the superposition of the flexibility profiles of collagen IV in sodium acetate and TBS buffers, including the respective 95% confidence intervals of the effective persistence lengths. Over the vast majority of the collagenous domain, the effective persistence length is statistically unaffected by this change in solution conditions upon deposition. This contrasts with the strong mechanical response of collagens to decreases in ionic strength (10). Over the range of pH from ~3 to 7.4, the net charge on collagen IV is expected to change significantly, from positively charged to approximately neutral (Fig. S13). Despite this change in charge, the net persistence length of collagen IV remains the same (*p* = 40 nm), and its position-dependent flexibility is essentially invariant (Fig. S14). This finding provides further evidence that the conformations we observe are not governed by electrostatic interactions with the surface. Furthermore, it implies that electrostatic interactions such as salt bridges (100) do not contribute substantially to the net bending stiffness of collagen at physiologically relevant ionic strengths.

There are only a few regions in the effective persistence length profile where the flexibility differs between these two buffered solution conditions. The region ~170 nm from the NC1 domain is of interest as a potential site for NC1 binding that may lead to higher-order assembly (36). Sensitivity of the structure and mechanics in this region of the collagenous domain to solution pH and Cl^-^ could provide a responsive element that prevents premature assembly inside the cell. AFM-based mapping of ligand binding sites along collagen IV could help to reveal how these interactions are modulated by pH and chemical environment, and to relate the binding to local collagen structure. Sequence-based predictions of binding sites on collagen IV have been made for various binding partners (35, 101), which could be compared with AFM-based mapping experiments. Of particular interest would be proteins implicated in chaperoning collagen folding (e.g. Hsp47), secretion (e.g. TANGO1), and assembly (e.g. SPARC), as well as other proteins involved in basement membrane assembly (102, 103). For example, EM-based mapping has found laminin, a major BM component, to bind 140 nm from the NC1 domain along the collagen IV molecule (37). Analysis of our persistence length profile shows that this binding site is at a flexible region, which coincides with a large overlapping interruption. On the other hand, the glycoprotein nidogen was found to bind 80 nm from the NC1 domain (41), within a triple-helical region. These results suggest that there may be a broad diversity of binding modes to collagen IV, which can be modulated by its local structure and perhaps further tuned by the chemical environment. Studies of the temperature-dependent flexibility profile, binding interactions, and *in vitro* assembly morphologies could elucidate the impact of sequence-dependent mechanical and thermal stability on guiding BM assembly.

Our findings on the bending flexibility of collagen can be used to quantify energetic costs of various candidate compartments and constraints along its secretory and assembly pathways. For example, the enhanced flexibility of collagen IV imparted by its interruptions means that it is expected to adopt more compact conformations in solution compared with its shorter fibril-forming collagen counterparts. Thus, its secretory burden on the cell machinery may be less. However, this flexibility of collagen IV endows a greater entropic cost for lateral ordering into higher-order structures, compared with the stiffer collagen III. The greater rigidity of continuously triple-helical collagens facilitates their lateral assembly into highly ordered thick fibrils. We speculate that these mechanical properties contributed to the development of collagen in evolution, in which collagen IV arose first, but now is a minority component – at least by mass portion – compared with fibril-forming collagens that form the basis of most connective tissues in higher-level organisms (6).

## Conclusions

AFM-based mapping offers the ability to measure local structural properties such as bending stiffness within the context of full-length collagen proteins, and correlate these with sequence-dependent properties elucidated from studies on much shorter collagen-mimicking peptides. Our results reveal a heterogeneous mechanical response along the triple-helical domains of collagens. We found that the presence of interruptions provides significant flexibility to collagen IV, and were able to describe the shape of its flexibility profile reasonably well with a simple physical model that classifies sequences solely based on the presence of interruptions in the (Gly-X-Y) pattern. In our complementary imaging and analysis of collagen III molecules, we found significant mechanical heterogeneity along their lengths, indicating that a more refined picture of collagen flexibility should include sequence-dependent flexibility within triple-helical regions. Notably, we found collagen III to possess a unique mechanical signature at the MMP site, a key sequence for collagen’s extracellular remodelling. Surprisingly – in light of past suggestions that proline content influences local triple helix rigidity (17, 66, 67, 69, 70) – we find no net correlation between local proline abundance and flexibility of collagenous domains, though some tracts in both collagens III and IV exhibit an anticorrelation between flexibility and imino acid content. In this study, our analysis was limited to determining effective persistence lengths over 30-nm regions and deconvolving the effect of local sequence; future improvements in imaging and image analysis that allow more refined backbone tracing and registry of chains should further enhance our understanding of how local sequence influences mechanical properties of collagens.

## Supporting information

Sequence alignments

## Data availability

Primary and analysed data are available from the corresponding author upon request. Sequence alignments used in this work are provided in a Supporting document.

## Acknowledgements

We gratefully acknowledge Albert Ries for the gift of collagen IV from the Klaus Kühn lab, and Takako Sasaki for help procuring the sample. We thank Hans Peter Bächinger for providing the pN-III collagen and for supportive feedback. We thank Mike Kirkness, Mathew Schneider, Daniel Sloseris and other members of the Forde lab for useful suggestions on this work, and ChangMin Kim for assistance with AFM instrumentation.

## Author contributions

AAS performed all experiments and analysed all experimental data. AL devised and validated the position-dependent analysis protocols. AAS and NRF developed the variable flexibility model. All authors contributed to the project design and to writing the manuscript.

## Supporting Information available online

Supporting Information document: Includes Supporting Texts 1, 2; Table S1; Figures S1-S14.

Sequence alignments: Document providing different sequence alignments tested in this work.

Supporting citations: References (104–106) appear in the supporting material.

## References

1. Shoulders, M.D. and Raines, R.T. (2009) Collagen Structure and Stability. Annu. Rev. Biochem., 78, 929–958.

2. Sorushanova, A., Delgado, L.M., Wu, Z., Shologu, N., Kshirsagar, A., Raghunath, R., Mullen, A.M., Bayon, Y., Pandit, A., Raghunath, M., et al. (2019) The Collagen Suprafamily: From Biosynthesis to Advanced Biomaterial Development. Adv. Mater., 31, 1–39.

3. Fidler, A.L., Boudko, S.P., Rokas, A. and Hudson, B.G. (2018) The triple helix of collagens – an ancient protein structure that enabled animal multicellularity and tissue evolution. J. Cell Sci., 131.

4. Timpl, R., Wiedemann, H., Van Delden, V., Furthmayer, H. and Kühn, K. (1981) A Network Model for the Organization of Type IV Collagen Molecules in Basement Membranes. Eur. J. Biochem., 120, 203–211.

5. Knupp, C. and Squire, J.M. (2005) Molecular packing in network-forming collagens. Adv. Protein Chem., 70, 375–403.

6. Fidler, A.L., Darris, C.E., Chetyrkin, S. V, Pedchenko, V.K., Boudko, S.P., Brown, K.L., Gray Jerome, W., Hudson, J.K., Rokas, A. and Hudson, B.G. (2017) Collagen IV and basement membrane at the evolutionary dawn of metazoan tissues. Elife, 6.

7. Hudson, B.G. (2004) The molecular basis of goodpasture and alport syndromes: Beacons for the discovery of the collagen IV family. J. Am. Soc. Nephrol., 15, 2514–2527.

8. Kirkness, M.W., Lehmann, K. and Forde, N.R. (2019) Mechanics and structural stability of the collagen triple helix. Curr. Opin. Chem. Biol., 53, 98–105.

9. Fratzl, P. (2008) Collagen: Structure and Mechanics.

10. Rezaei, N., Lyons, A. and Forde, N.R. (2018) Environmentally controlled curvature of single collagen proteins. Biophys. J., 115, 1457–1469.

11. Miller, R.T. (2017) Mechanical properties of basement membrane in health and disease. Matrix Biol., 57-58, 366–373.

12. Kadler, K.E., Hojima, Y. and Prockop, D.J. (1987) Assembly of Collagen Fibrils de Novo by cleavage of the Type 1 pC-Collagen with Procollagen C-Proteinase. J. Biol. Chem., 260, 15696–15701.

13. Brown, K.L., Cummings, C.F., Vanacore, R.M. and Hudson, B.G. (2017) Building collagen IV smart scaffolds on the outside of cells. Protein Sci., 26, 2151–2161.

14. Cummings, C.F., Pedchenko, V., Brown, K.L., Colon, S., Rafi, M., Jones-Paris, C., Pokydeshava, E., Liu, M., Pastor-Pareja, J.C., Stothers, C., et al. (2016) Extracellular chloride signals collagen IV network assembly during basement membrane formation. J. Cell Biol., 213, 479–494.

15. Hwang, E.S. and Brodsky, B. (2012) Folding delay and structural perturbations caused by type IV collagen natural interruptions and nearby Gly missense mutations. J. Biol. Chem., 287, 4368–4375.

16. Bella, J. (2014) A first census of collagen interruptions: Collagen’s own stutters and stammers. J. Struct. Biol., 186, 438–450.

17. Hofmann, H., Voss, T. and Kühn, K. (1984) Localization of Flexible Sites in Thread-like Molecules from Electron Micrographs - Comparison of Interstitial, Basement Membrane and Intima Collagens. J. Mol. Biol., 172, 325–343.

18. Lunstrum, G.P., Bächinger, H.P., Fessler, L.I., Duncan, K.G., Nelson, R.E. and Fessler, J.H. (1988) Drosophila basement membrane procollagen IV. I. Protein characterization and distribution. J. Biol. Chem., 263, 18318–18327.

19. Bächinger, H.P., Doege, K.J., Petschek, J.P., Fessler, L.I. and Fessler, J.H. (1982) Structural Implications from an Electronmicroscopic Comparison of Procollagen V with Procollagen I, pC-Collagen I, Procollagen IV, and a Drosophila Procollagen. J. Biol. Chem., 257, 14590–14592.

20. Bächinger, H.P., Morris, N.P., Lunstrums, G.P., Keene, D.R., Rosenbaum, L.M., Compton, L.A. and Burgeson, R.E. (1990) The Relationship of the Biophysical and Biochemical Characteristics of Type VII Collagen to the Function of Anchoring Fibrils. J. Biol. Chem., 265, 10095–10101.

21. Thiagarajan, G., Li, Y., Mohs, A., Strafaci, C., Popiel, M., Baum, J. and Brodsky, B. (2008) Common Interruptions in the Repeating Tripeptide Sequence of Non-fibrillar Collagens: Sequence Analysis and Structural Studies on Triple-helix Peptide Models. J. Mol. Biol., 376, 736–748.

22. Hwang, E.S., Thiagarajan, G., Parmar, A.S. and Brodsky, B. (2010) Interruptions in the collagen repeating tripeptide pattern can promote supramolecular association. Protein Sci., 19, 1053–1064.

23. Bella, J. (2016) Collagen structure: new tricks from a very old dog. Biochem. J., 473, 1001–1025.

24. Orgel, J.P.R.O., Irving, T.C., Miller, A. and Wess, T.J. (2006) Microfibrillar structure of type I collagen in situ. Proc. Natl. Acad. Sci. U. S. A., 103, 9001–9005.

25. Antipova, O. and Orgel, J.P.R.O. (2010) In situ D-periodic molecular structure of type II collagen. J. Biol. Chem., 285, 7087–7096.

26. Bella, J., Eaton, M., Brodsky, B. and Berman, H.M. (1994) Crystal and molecular structure, of a collagen-like peptide at 1.9 Å resolution. Science (80-.)., 266, 75–81.

27. Golbik, R., Eble, J.A., Ries, A. and Kühn, K. (2000) The spatial orientation of the essential amino acid residues arginine and aspartate within the α1β1 integrin recognition site of collagen IV has been resolved using fluorescence resonance energy transfer. J. Mol. Biol., 297, 501–509.

28. Kleinman, H.K., McGarvey, M.L., Liotta, L.A., Robey, P.G., Tryggvason, K. and Martin, G.R. (1982) Isolation and Characterization of Type IV Procollagen, Laminin, and Heparan Sulfate Proteoglycan from the EHS Sarcoma. Biochemistry, 21, 6188–6193.

29. Vandenberg, P., Kern, A., Ries, A., Luckenbill-edds, L., Mann, K. and Kühn, K. (2000) Characterization of a type IV collagen major cell binding site with affinity to the α1/β1 and the α2/β1 integrins. J. Cell Biol., 113, 1475–83.

30. Timpl, R., Glanville, R.W., Nowack, H., Wiedemann, H., Fietzek, P.P. and Kühn, K. (1975) Isolation, Chemical and Electron Microscopical Characterization of Neutral-Salt-Soluble Type III Collagen and Procollagen from Fetal Bovine Skin. Hoppe-Seyler’s Zeitschrift für Physiol. Chemie, 56, 1783–1792.

31. Schneider, M., Al-Shaer, A. and Forde, N. (2021) AutoSmarTrace: Automated AutoSmarTrace: Automated Chain Tracing and Flexibility Analysis of Biological Filaments. bioRxiv, https://doi.org/10.1101/2021.01.19.427319.

32. MATLAB and Statistical Toolbox Release 2018b. The MathWorks, Inc., Natick, Massachusetts, United States.

33. Bateman, A. (2019) UniProt: A worldwide hub of protein knowledge. Nucleic Acids Res., 47, D506–D515.

34. Kühn, K. (1995) Basement membrane (type IV) collagen. Matrix Biol., 14, 439–445.

35. Parkin, J. Des, Antonio, J.D.S., Pedchenko, V., Hudson, B., Jensen, S.T. and Savige, J. (2011) Mapping Structural Landmarks, Ligand Binding Sites, and Missense Mutations to the Collagen IV Heterotrimers Predicts Major Functional Domains, Novel Interactions, and Variation in Phenotypes in Inherited Diseases Affecting Basement Membranes. Hum. Mutat., 32, 127–143.

36. Yurchenco, P.D. and Furthmayr, H. (1984) Self-Assembly of Basement Membrane Collagen. Biochemistry, 23, 1839–1850.

37. Charonis, A.S., Tsilibary, E.C., Yurchenco, P.D., Furthmayr, H. and Coritz, A. (1985) Binding of laminin to type IV collagen: A morphological study. J. Cell Biol., 100, 1848–1853.

38. Laurie, G.W., Bing, J.T., Kleinman, H.K., Hassell, J.R., Aumailley, M., Martin, G.R. and Feldmann, R.J. (1986) Localization of binding sites for laminin, heparan sulfate proteoglycan and fibronectin on basement membrane (type IV) collagen. J. Mol. Biol., 189, 205–216.

39. Tsilibary, E.C. and Charonis, A.S. (1986) The role of the main noncollagenous domain (NC1) in type IV collagen self-assembly. J. Cell Biol., 103, 2467–2473.

40. Yurchenco, P.D. and Ruben, G.C. (1987) Basement-Membrane Structure Insitu - Evidence for Lateral Associations in the Type-Iv Collagen Network. J. Cell Biol., 105, 2559–2568.

41. Aumailley, M., Wiedemann, H., Mann, K. and Timpl, R. (1989) Binding of nidogen and the laminin-nidogen complex to basement membrane collagen type IV. Eur. J. Biochem., 184, 241–248.

42. Fox, J.W., Mayer, U., Nischt, R., Aumailley, M., Reinhardt, D., Wiedemann, H., Mann, K., Timpl, R., Krieg, T. and Engel, J. (1991) Recombinant nidogen consists of three globular domains and mediates binding of laminin to collagen type IV. EMBO J., 10, 3137–3146.

43. Lovelady, H.H., Shashidhara, S. and Matthews, W.G. (2014) Solvent specific persistence length of molecular type I collagen. Biopolymers, 101, 329–335.

44. Pedchenko, V., Bauer, R., Pokidysheva, E.N., Al-Shaer, A., Forde, N.R., Fidler, A.L., Hudson, B.G. and Boudko, S.P. (2019) A chloride ring is an ancient evolutionary innovation mediating the assembly of the collagen IV scaffold of basement membranes. J. Biol. Chem., 294, jbc.RA119.007426.

45. Pedchenko, V., Boudko, S.P., Barber, M., Mikhailova, T., Saus, J., Harmange, J.-C. and Hudson, B.G. (2021) Collagen IVα345 dysfunction in glomerular basement membrane diseases. III. A functional framework for α345 hexamer assembly. J. Biol. Chem., 296, 100592.

46. Chen, C.H. and Hansma, H.G. (2000) Basement membrane macromolecules: Insights from atomic force microscopy. J. Struct. Biol., 131, 44–55.

47. Rivetti, C., Guthold, M. and Bustamante, C. (1996) Scanning force microscopy of DNA deposited onto mica: Equilibration versus kinetic trapping studied by statistical polymer chain analysis. J. Mol. Biol., 264, 919–932.

48. Heenan, P.R. and Perkins, T.T. (2019) Imaging DNA Equilibrated onto Mica in Liquid Using Biochemically Relevant Deposition Conditions. ACS Nano, 13, 4220–4229.

49. Davis, J.M., Boswell, B.A. and Bächinger, H.P. (1989) Thermal stability and folding of type IV procollagen and effect of peptidyl-prolyl cis-trans-isomerase on the folding of the triple helix. J. Biol. Chem., 264, 8956–8962.

50. Shayegan, M., Altindal, T., Kiefl, E. and Forde, N.R. (2016) Intact Telopeptides Enhance Interactions between Collagens. Biophysj, 111, 2404–2416.

51. Li, H., Linke, W.A., Oberhauser, A.F., Carrion-Vazquez, M., Kerkvliet, J.G., Lu, H., Marszalek, P.E. and Fernandez, J.M. (2002) Reverse engineering of the giant muscle protein titin. Nature, 418, 998–1002.

52. Rivetti, C., Walker, C. and Bustamante, C. (1998) Polymer chain statistics and conformational analysis of DNA molecules with bends or sections of different flexibility. J. Mol. Biol., 280, 41–59.

53. Todd, B.A., Rammohan, J. and Eppell, S.J. (2003) Connecting nanoscale images of proteins with their genetic sequences. Biophys. J., 84, 3982–3991.

54. Valdman, D., Lopez, B.J., Valentine, M.T. and Atzberger, P.J. (2013) Force spectroscopy of complex biopolymers with heterogeneous elasticity. Soft Matter, 9, 772–778.

55. Brazel, D., Pollner, R., Oberbaumer, I. and Kühn, K. (1988) Human basement membrane collagen (type IV) The amino acid sequence of the a2(IV) chain and its comparison with the al(1V) chain reveals deletions in the al(1V) chain. Eur J Biochem, 172, 35–42.

56. Engel, J. and Prockop, D.J. (1991) The Zipper-Like Folding of Collagen Triple Helices and the Effects of Mutations that Disrupt the Zipper. Annu. Rev. Biophys. Biophys. Chem., 20, 137–152.

57. Orgel, J.P.R.O., Persikov, A. V. and Antipova, O. (2014) Variation in the helical structure of native collagen. PLoS One, 9.

58. Bann, J.G., Peyton, D.H. and Bächinger, H.P. (2000) Sweet is stable: Glycosylation stabilizes collagen. FEBS Lett., 472, 237–240.

59. Perdivara, I., Yamauchi, M. and Tomer, K.B. (2013) Molecular characterization of collagen hydroxylysine O-glycosylation by mass spectrometry: Current status. Aust. J. Chem., 66, 760–769.

60. Tang, M., Wang, X., Gandhi, N.S., Foley, B.L., Burrage, K., Woods, R.J. and Gu, Y. (2020) Effect of hydroxylysine-O-glycosylation on the structure of type I collagen molecule: A computational study. Glycobiology, 00, 1–14.

61. Basak, T., Vega-Montoto, L., Zimmerman, L.J., Tabb, D.L., Hudson, B.G. and Vanacore, R.M. (2016) Comprehensive Characterization of Glycosylation and Hydroxylation of Basement Membrane Collagen IV by High-Resolution Mass Spectrometry. J. Proteome Res., 15, 245–258.

62. Liotta, L.A., Abe, S., Gehron Robey, P. and Martin, G.R. (1979) Preferential digestion of basement membrane collagen by an enzyme derived from a metastatic murine tumor. Proc. Natl. Acad. Sci. U. S. A., 76, 2268–2272.

63. Srinivasan, M., Uzel, S.G.M., Gautieri, A., Keten, S. and Buehler, M.J. (2009) Alport Syndrome mutations in type IV tropocollagen alter molecular structure and nanomechanical properties. J. Struct. Biol., 168, 503–510.

64. Wu, Y.Y., Bao, L., Zhang, X. and Tan, Z.J. (2015) Flexibility of short DNA helices with finite-length effect: From base pairs to tens of base pairs. J. Chem. Phys., 142.

65. Bächinger, H.P., Bruckner, P., Timpl, R., Prockop, D.J. and Engel, J. (1980) Folding Mechanism of the Triple Helix in Type-III Collagen and Type-III pN–Collagen: Role of Disulfide Bridges and Peptide Bond Isomerization. Eur. J. Biochem., 106, 619–632.

66. Holmes, D.F., Mould, A.P. and Chapman, J.A. (1991) Morphology of sheet-like assemblies of pN-collagen, pC-collagen and procollagen studied by scanning transmission electron microscopy mass measurements. J. Mol. Biol., 10.1016/0022-2836(91)90385-J.

67. Bruckner, P., Bächinger, H.P., Timpl, R. and Engel, J. (1978) Three Conformationally Distinct Domains in the Amino-Terminal Segment of Type III Procollagen and Its Rapid Triple Helix + Coil Transition. Eur. J. Biochem., 603, 595–603.

68. Bhatnagar, R.S., Qian, J.J. and Gough, C.A. (1997) The role in cell binding of a β1-bend within the triple helical region in collagen α1(i) chain: Structural and biological evidence for conformational tautomerism on fiber surface. J. Biomol. Struct. Dyn., 14, 547–560.

69. Farndale, R.W., Lisman, T., Bihan, D., Hamaia, S., Smerling, C.S., Pugh, N., Konitsiotis, A., Leitinger, B., De Groot, P.G., Jarvis, G.E., et al. (2008) Cell-collagen interactions: The use of peptide Toolkits to investigate collagen-receptor interactions. Biochem. Soc. Trans., 36, 241–250.

70. Lisman, T., Raynal, N., Groeneveld, D., Maddox, B., Peachey, A.R., Huizinga, E.G., De Groot, P.G. and Farndale, R.W. (2006) A single high-affinity binding site for von Willebrand factor in collagen III, identified using synthetic triple-helical peptides. Blood, 108, 3753–3756.

71. Lauer-Fields, J.L., Juska, D. and Fields, G.B. (2002) Matrix metalloproteinases and collagen catabolism. Biopolym. - Pept. Sci. Sect., 66, 19–32.

72. Ravikumar, K.M. and Hwang, W. (2008) Region-specific role of water in collagen unwinding and assembly. Proteins, 72, 1320–1332.

73. Miller, E.J., Finch, J.E., Chung, E., Butler, W.T. and Robertson, P.B. (1976) Specific cleavage of the native Type III collagen molecule with trypsin. Similarity of the cleavage products to collagenase-produced fragments and primary structure at the cleavage site. Arch. Biochem. Biophys., 173, 631–637.

74. Williams, K.E. and Olsen, D.R. (2009) Matrix metalloproteinase-1 cleavage site recognition and binding in full-length human type III collagen. Matrix Biol., 28, 373–379.

75. Kirkness, M.W.H. and Forde, N.R. (2018) Single-Molecule Assay for Proteolytic Susceptibility: Force-Induced Collagen Destabilization. Biophys. J., 114, 570–576.

76. Fields, G.B. (1991) A model for interstitial collagen catabolism by mammalian collagenases. J. Theor. Biol., 153, 585–602.

77. Stultz, C.M. (2002) Localized unfolding of collagen explains collagenase cleavage near imino-poor sites. J. Mol. Biol., 319, 997–1003.

78. Glanville, R.W., Voss, T. and Kühn, K. (1982) A Comparison of the Flexibility of Molecules of Basement Membrane and Interstitial Collagens.

79. Chow, W.Y., Forman, C.J., Bihan, D., Puszkarska, A., Rajan, R., Reid, D.G., Colwell, L.J., Wales, D.J., Farndale, R.W. and Melinda J. Duer (2018) Proline provides site-specific flexibility for in vivo collagen. Sci. Rep., 8, 1–13.

80. Weis, M.A., Hudson, D.M., Kim, L., Scott, M., Wu, J.J. and Eyre, D.R. (2010) Location of 3-Hydroxyproline residues in collagen types I, II, III, and V/XI implies a role in fibril supramolecular assembly. J. Biol. Chem., 285, 2580–2590.

81. Montgomery, N.T., Zientek, K.D., Pokidysheva, E.N. and Bächinger, H.P. (2018) Post-translational modification of type IV collagen with 3-hydroxyproline affects its interactions with glycoprotein VI and nidogens 1 and 2. J. Biol. Chem., 293, 5987–5999.

82. Taga, Y., Tanaka, K., Hattori, S. and Mizuno, K. (2021) In-depth correlation analysis demonstrates that 4-hydroxyproline at the Yaa position of Gly-Xaa-Yaa repeats dominantly stabilizes collagen triple helix. Matrix Biol. Plus, 10.1016/j.mbplus.2021.100067.

83. Berg, R.A. and Prockop, D.J. (1973) The thermal transition of a non-hydroxylated form of collagen. Evidence for a role for hydroxyproline in stabilizing the triple-helix of collagen. Biochem. Biophys. Res. Commun., 52, 115–120.

84. Burjanadze, T. V. (1979) Hydroxyproline content and location in relation to collagen thermal stability. Biopolymers, 18, 931–938.

85. Uzel, S.G.M. and Buehler, M.J. (2009) Nanomechanical sequencing of collagen: Tropocollagen features heterogeneous elastic properties at the nanoscale. Integr. Biol., 1, 452–459.

86. Gautieri, A., Vesentini, S., Redaelli, A. and Buehler, M.J. (2009) Single molecule effects of osteogenesis imperfecta mutations in tropocollagen protein domains. Protein Sci., 18, 161–168.

87. Yeo, J., Qiu, Y., Jung, G.S., Zhang, Y.W., Buehler, M.J. and Kaplan, D.L. (2020) Adverse effects of Alport syndrome-related Gly missense mutations on collagen type IV: Insights from molecular simulations and experiments. Biomaterials, 240, 119857.

88. Malhotra, V. and Erlmann, P. (2015) The Pathway of Collagen Secretion. Annu. Rev. Cell Dev. Biol., 31, 109–124.

89. Stauber, T. and Jentsch, T.J. (2013) Chloride in Vesicular Trafficking and Function. Annu. Rev. Physiol., 75, 453–477.

90. Dölz, R., Engel, J. and Kühn, K. (1988) Folding of collagen IV. Eur. J. Biochem., 178, 357–366.

91. Weber, S., Engel, J., Wiedemann, H., Glanville, R.W. and Timpl, R. (1984) Subunit structure and assembly of the globular domain of basement-membrane collagen type IV. Eur. J. Biochem., 139, 401–410.

92. Wood, G.C. and Keech, M.K. (1960) The formation of fibrils from collagen solutions 1. The effect of experimental conditions: kinetic and electron-microscope studies. Biochem. J., 75, 588–598.

93. Harris, J.R., Soliakov, A. and Lewis, R.J. (2013) In vitro fibrillogenesis of collagen type I in varying ionic and pH conditions. Micron, 49, 60–68.

94. Roberto Vanacore, Amy-Joan L. Ham, Markus Voehler, Charles R. Sanders, Thomas P. Conrads, Timothy D. Veenstra, K. Barry Sharpless, Philip E. Dawson, B.G.H. (2009) A Sulfilimine Bond Identified in Collagen IV. Science, 325, 1230–1234.

95. McCaughey, J. and Stephens, D.J. (2019) ER-to-Golgi Transport: A Sizeable Problem. Trends Cell Biol., 29, 940–953.

96. Raote, I., Ortega-Bellido, M., Santos, A.J.M., Foresti, O., Zhang, C., Garcia-Parajo, M.F., Campelo, F. and Malhotra, V. (2018) TANGO1 builds a machine for collagen export by recruiting and spatially organizing COPII, tethers and membranes. Elife, 10.7554/eLife.32723.

97. Matsui, Y., Hirata, Y., Wada, I. and Hosokawa, N. (2020) Visualization of procollagen IV reveals ER-to-Golgi transport by ERGIC-independent carriers. Cell Struct. Funct., 10.1247/csf.20025.

98. McCaughey, J., Stevenson, N.L., Cross, S. and Stephens, D.J. (2019) ER-to-Golgi trafficking of procollagen in the absence of large carriers. J. Cell Biol., 218, 929–948.

99. Freudenberg, U., Behrens, S.H., Welzel, P.B., Müller, M., Grimmer, M., Salchert, K., Taeger, T., Schmidt, K., Pompe, W. and Werner, C. (2007) Electrostatic interactions modulate the conformation of collagen I. Biophys. J., 92, 2108–2119.

100. Keshwani, N., Banerjee, S., Brodsky, B. and Makhatadze, G.I. (2013) The role of cross-chain ionic interactions for the stability of collagen model peptides. Biophys. J., 105, 1681–1688.

101. Hohenester, E., Sasaki, T., Giudici, C., Farndale, R.W. and Bächinger, H.P. (2008) Structural basis of sequence-specific collagen recognition by SPARC. Proc. Natl. Acad. Sci. U. S. A., 105, 18273–18277.

102. Chioran, A., Duncan, S., Catalano, A., Brown, T.J. and Ringuette, M.J. (2017) Collagen IV trafficking: The inside-out and beyond story. Dev. Biol., 431, 124–133.

103. Köhler, A., Mörgelin, M., Gebauer, J.M., Öcal, S., Imhof, T., Koch, M., Nagata, K., Paulsson, M., Aumailley, M., Baumann, U., et al. (2020) New specific HSP47 functions in collagen subfamily chaperoning. FASEB J., 10.1096/fj.202000570R.

104. Landau, L. D., L. P. Pitaevskii, A. M. Kosevich, and E.M.L. (1986) Theory of Elasticity. Butterworth-Heinemann.

105. Krishnamoorthy, K. (2016) Handbook of statistical distribution with applications (2nd edition). CRC Press: Boca Raton, Florida.

106. Gelman, A. (2013) Bayesian Data Analysis (3rd edition). CRC Press: Boca Raton, Florida.

