## Supporting Information for "Sequence-dependent mechanics of collagen reflect its structural and functional organization"

**Contents**

**Supporting Text 1**

Inhomogeneous Worm-like Chain Theory and Application……….………….………………2

**Figure S1**. Differences among theoretical persistence length profiles...……….……......3

Statistical Properties of Persistence Length Estimates…………………………………....4

**Figure S2**. Probability density of persistence length estimates, for *p* = 85 nm and *n* = 100 chains……………..……………………………...…………………………………..…5

**Figure S3**. Dependence of persistence length estimate and 95% confidence interval on sample size ……………….……...…………………………….…………………….……....6

**Figure S4.** Effective persistence length vs. segment length…………………………..………..7

**Figure S5**. Amino acid sequences of the collagen IV collagenous domain…………„…....…..8

**Figure S6.** Flexibility profiles of collagen IV traced in both directions.…………………….…10

**Figure S7**. Varying assumptions of the minimum number of tripeptide units required to form triple-helical segments………......………….……………………………………..…...……..11

**Supporting Text 2**

Variable Flexibility Model Fitting………………………………………………..……..12

**Table S1:** Model outputs of flexibility for different chain alignments……………..…..14

**Figure S8.** Four-class flexibility model using a linear chain alignment………………..…..…...15

**Figure S9.** Flexibility profile of collagen pN-III traced from the N terminus...……………..…16

**Figure S10**. Proline content and persistence length profile of pN-III collagen………………....17

**Figure S11**. Proline content and persistence length profile of collagen IV…………………..18

**Figure S12**. Homogeneous worm-like chain determination of persistence lengths of collagen IV in different solution conditions……………………………………………….……….......19

**Figure S13.** Charge profiles of the α1 and α2 chains of collagen IV at different pH……..........20

**Figure S14**. Superposition of experimental persistence length profiles in all three solution conditions……………………………………………………………..………..……..22

**Supporting References**…………………………………………………………………….23

### Supporting Text 1

#### Inhomogeneous Worm-like Chain Theory and Application

Consider a section of a two-dimensional worm-like chain with length $\Delta s$, broken up into $n$ infinitesimal segments of length $\delta s = \frac{\Delta s}{n}$. The energy required to bend segment $i$ into a circular arc is given by

$$E_i = \frac{\alpha_i \delta\theta_i^2}{2\delta s} = \frac{p_i \delta\theta_i^2}{2\delta s} k_\mathrm{B}T. \quad \text{(S1)}$$

$\alpha_i$ is the bending rigidity of the segment, related to the persistence length $p_i$ through $p_i = \alpha_i / k_\mathrm{B}T$, where $k_\mathrm{B}T$ is the product of the Boltzmann constant and the absolute temperature (1). The central angle of this arc, $\delta\theta_i$, is equivalent to the angle between the tangents at the beginning and end of the segment. At equilibrium, the distribution of angles $\delta\theta_i$ is given by the Boltzmann distribution, where

$$P(\delta\theta_i) = \sqrt{\frac{p_i}{2\pi\delta s}} \exp\left(\frac{p_i \delta\theta_i^2}{2\delta s}\right). \quad \text{(S2)}$$

This is a normal distribution with $\langle\delta\theta_i\rangle = 0$ and variance $\sigma_{\delta\theta_i}^2 = \frac{\delta s}{p_i}$.

Since each $\delta\theta_i$ is a signed angle, the total angle adopted over the total section length $\Delta s$ is given by $\theta = \sum_{i=1}^{n} \delta\theta_i$. Using the fact that each $\delta\theta_i$ is governed by an independent normal distribution, the sum of each of these angles will also be normally distributed, and thus will have mean $\langle\theta\rangle = \sum_{i=1}^{n}\langle\delta\theta_i\rangle = 0$ and variance $\sigma_\theta^2 = \sum_{i=1}^{n} \sigma_{\delta\theta_i}^2 = \sum_{i=1}^{n} \frac{\delta s}{p_i}$. Taking the limit as $n \to \infty$, we therefore find that

$$\langle\theta\rangle = 0 \quad \text{(S3)}$$

and

$$\sigma_\theta^2 = \int_0^{\Delta s} \frac{ds'}{p(s')}. \quad \text{(S4)}$$

Here, $p(s')$ is the persistence length at position $0 \le s' \le \Delta s$ along the contour of the chain segment.

Equation (S4) has strong implications for the experimental determination of persistence length profiles: because estimates of persistence length require measurements over a finite segment of length $\Delta s$, variations in persistence length within this length cannot be determined. Instead, the angular variance measured across the segment is

$$\sigma_\theta^2 = \frac{\Delta s}{p^*}, \quad \text{(S5)}$$

where $p^*$ is the effective persistence length, taken to be uniform along the segment. Thus, for a given segment length $\Delta s$, the effective persistence length is calculated as

$$p^* = \frac{\Delta s}{\sigma_\theta^2} = \frac{\Delta s}{\int_0^{\Delta s} \frac{ds'}{p(s')}}. \quad \text{(S6)}$$

For polymers that have sharp changes in persistence length within their contours (e.g. collagen type IV, DNA with short single stranded regions), this effective persistence length will be biased towards the smaller values present in the measured segment. Consider the persistence length profile shown in Figure S1: the original persistence length profile $p(s)$ (shown in black) consists of five segments with 85 nm persistence lengths, interspaced with short, variable length sections with 5 nm persistence lengths. The effective persistence length profile (shown in red) calculated using Eq. (S6) is also plotted, using a measurement length of 30 nm. Lastly, for comparison purposes only, a simple average of the persistence length is plotted, calculated by averaging $p(s)$ over the same 30 nm window (shown in blue). We note that this average persistence length is *not* the correct way to view the effects of heterogeneous flexibility; rather, the effective persistence length (red) is expected, based on the angular flexibility of the 30-nm segments. In the main text, it is shown that the effective persistence length profile agrees well with that extracted from tracing images of simulated chains.

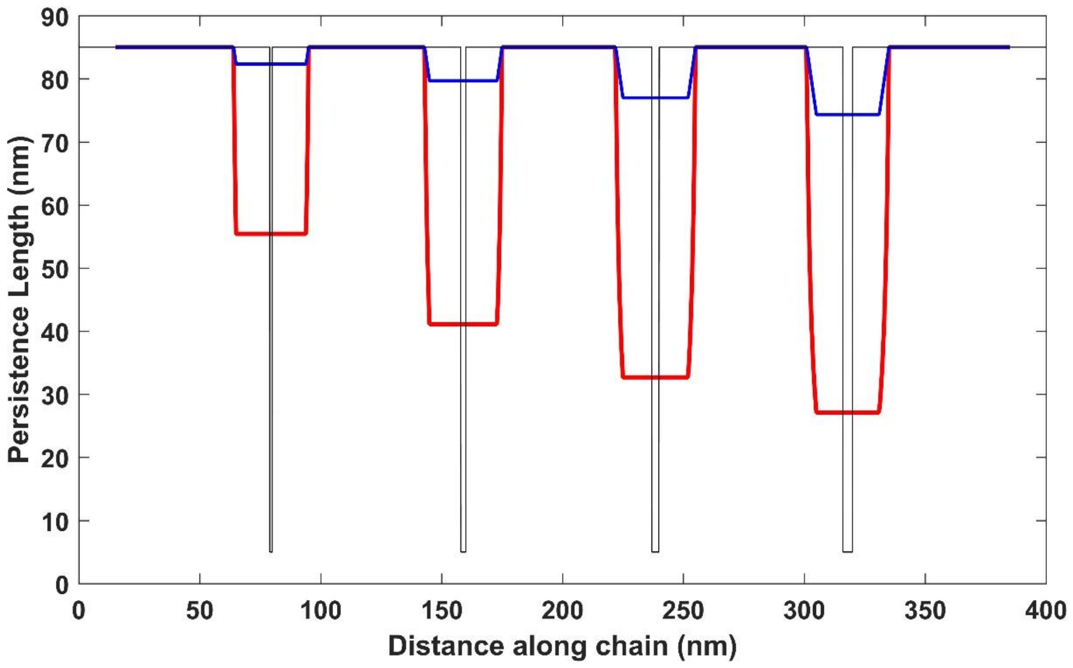


**Figure S1. Differences among theoretical persistence length profiles.** The original persistence length profile is shown in black, while the red line shows the effective persistence length profile calculated using Eq. (S6) and centered at the middle of the filter window. The average value of the original persistence length map over this 30 nm window is shown in blue for comparison.

**Statistical Properties of Persistence Length Estimates**

The persistence length is calculated using the variance of the angular distribution, which we can rewrite as follows, since $\langle\theta\rangle = 0$:

$$p = \frac{\Delta s}{\langle\theta^2\rangle}. \tag{S7}$$

For a finite number of observations, the quantity $\langle\theta^2\rangle$ is an estimate of the true variance, and can be represented by a random variable $S^2$ such that

$$S^2 = \frac{\Delta s}{p}\sum_{i=1}^{n}\frac{X_i^2}{n}, \tag{S8}$$

where $n$ is the number of observations and each random variable $X_i$ is drawn from a normal distribution with a mean of zero and variance of 1. $S^2$ can therefore be written in terms of the $\chi^2$ distribution as (2)

$$S^2 = \frac{\Delta s}{np}\chi_n^2. \tag{S9}$$

From this angular variance estimate, an estimate of the true value of the persistence length can be made. This estimate of $p$ is represented by a random variable, $P$, such that

$$P = \frac{\Delta s}{S^2} = \frac{\Delta s}{\left(\frac{\Delta s}{np}\chi_n^2\right)} = np\chi_n^{-2}. \tag{S10}$$

Here, $\chi_n^{-2}$ denotes the inverse chi-squared-distributed random variable. We can incorporate the $p$ and $n$ terms into this random variable, yielding

$$P \sim \text{Scale-Inv-}\chi^2(n, p), \tag{S11}$$

where Scale-Inv-$\chi^2(n, p)$ is a scaled inverse chi-squared-distributed random variable. This particular distribution has the probability density function (PDF) (3)

$$f(x; n, p) = \left(\frac{np}{2}\right)^{n/2}\frac{\exp\left(\frac{-pn}{2x}\right)}{x^{\frac{n}{2}+1}\Gamma\left(\frac{n}{2}\right)}, \tag{S12}$$

where $\Gamma\left(\frac{n}{2}\right) = \int_0^\infty t^{\frac{n}{2}-1}\mathrm{e}^{-t}\mathrm{d}t$ is the standard gamma function and $x$ represents the realization of $P$. The PDF described by equation (S12) is plotted in Figure S2 (in blue) for the case of $n = 100$ and $p = 85$ nm. The probability density is roughly centered around 85 nm (the actual persistence length), but estimates of the persistence length are more likely to be overestimated than underestimated.

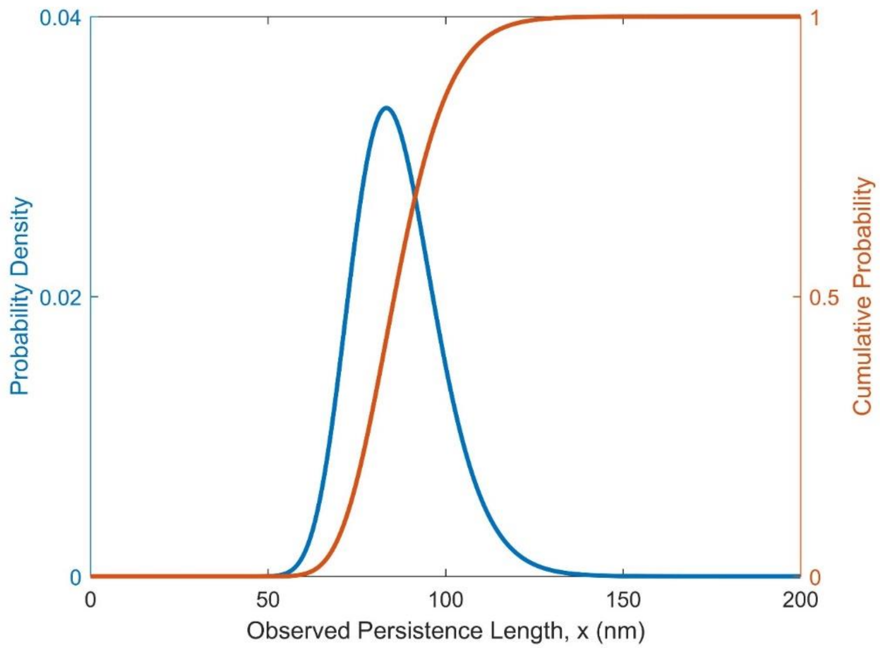


**Figure S2. Probability density of persistence length estimates, for *p* = 85 nm and *n* = 100 chains.** The blue line shows the expected distribution of persistence length estimates, obtained from equation (S12). The red line shows the cumulative probability of these estimates, given by equation (S13).

To calculate the confidence intervals on estimates of persistence length, we use the cumulative density function $F(x;n,p) = \int_0^x f(t;n,p)dt$. This function is given by

$$F(x;n,p) = \frac{1}{\Gamma\left(\frac{n}{2}\right)} \int_0^x \frac{1}{t}\left(\frac{pn}{2t}\right)^{n/2} e^{-\left(\frac{pn}{2t}\right)} dt = \frac{1}{\Gamma\left(\frac{n}{2}\right)} \int_{\frac{pn}{2x}}^{\infty} u^{\frac{n}{2}-1} e^{-u} du = \frac{\Gamma\left(\frac{n}{2},\frac{pn}{2x}\right)}{\Gamma\left(\frac{n}{2}\right)}, \quad \text{(S13)}$$

where the substitution $u = \frac{pn}{2t}$ was used to simplify the integral, and $\Gamma\left(\frac{n}{2},\frac{pn}{2x}\right) = \int_{\frac{pn}{2x}}^{\infty} t^{\frac{n}{2}-1} \mathrm{e}^{-t} \mathrm{d}t$ is the upper incomplete gamma function. This is plotted in red in Figure S2, for the same case of $n = 100$ and $p = 85$ nm.

Finally, we determine how the mean and 95% confidence intervals of the observed persistence length (the realization of $P$) depend on sample number. The mean value of the persistence length estimate is given by (3)

$$\langle P \rangle = p\left(\frac{n}{n-2}\right). \quad \text{(S14)}$$

This means that the observed persistence length will, on average, be overestimated relative to the true value for small $n$. To calculate the confidence intervals, the cumulative density function can be numerically inverted to find $x$ at $F(x;n,p) = 0.975$ and $F(x;n,p) = 0.025$ to extract the bounds on the 95% confidence interval. A plot of these values for $p = 85$ nm and $n = 100$ is shown in Figure S3.

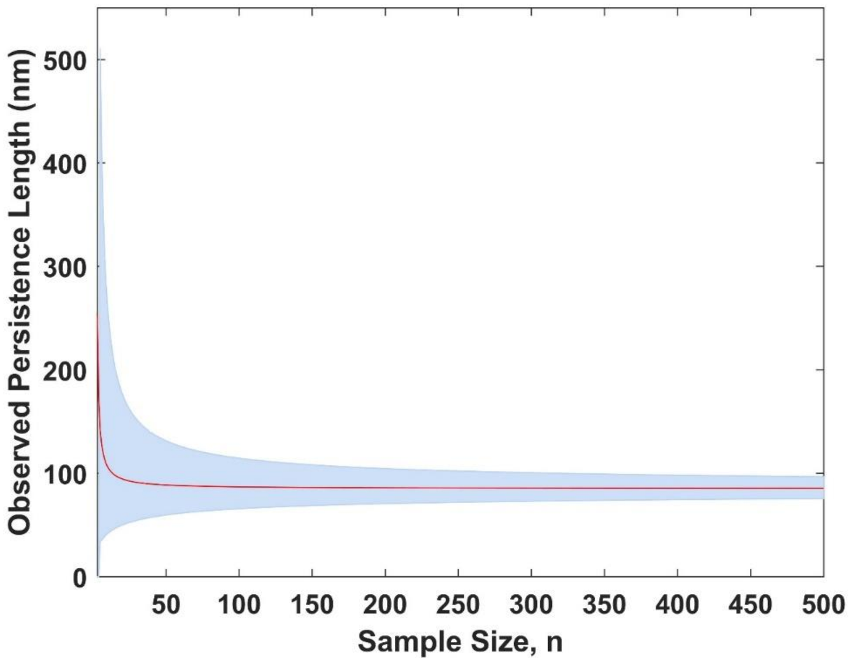


**Figure S3. Dependence of persistence length estimate and 95% confidence interval on sample size.** These values correspond to chains with persistence length $p$ = 85 nm. The mean value is obtained using equation (S14) and the error bounds are determined from the cumulative probability distribution equation (S13) as described in the text.

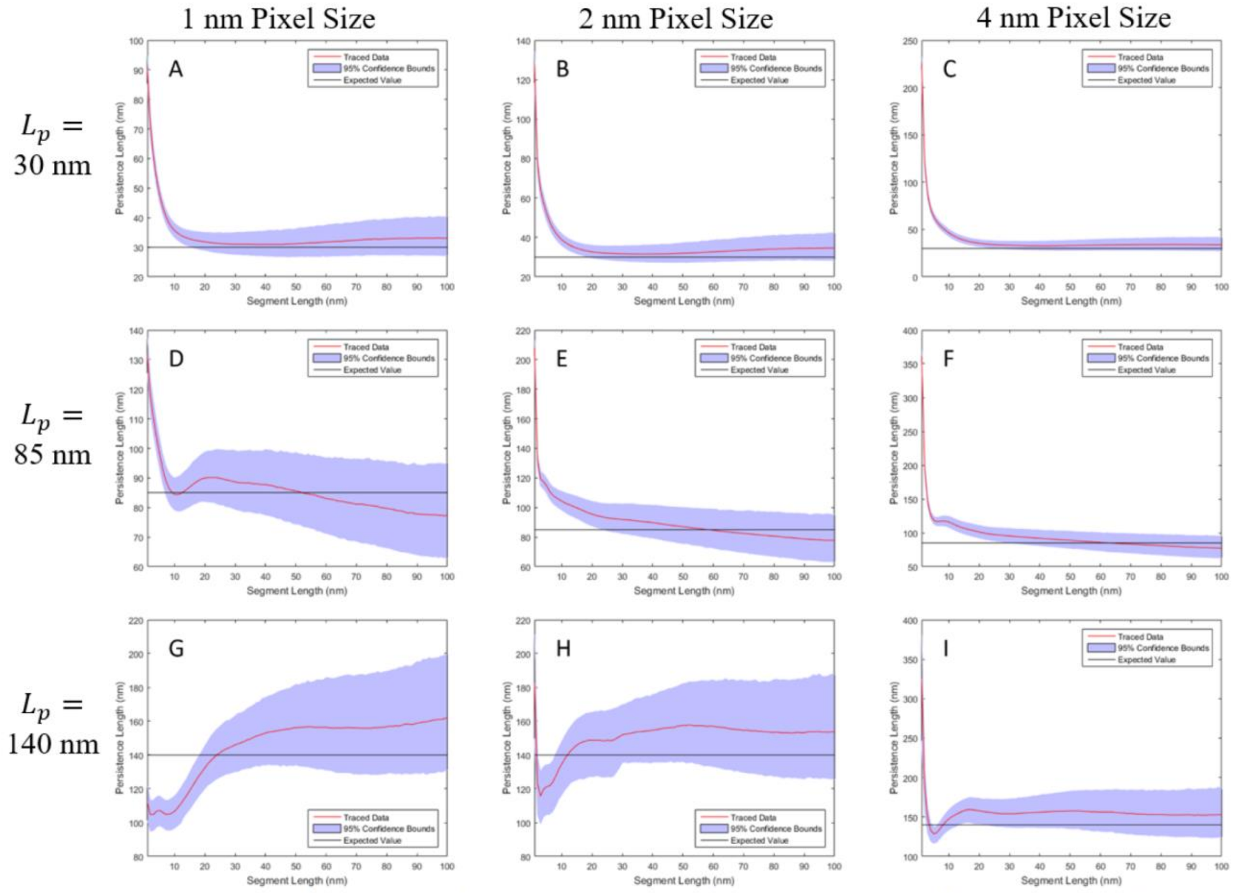


**Figure S4. Effective persistence length vs. segment length.** The effective persistence length $p^*$ was determined using Equation S5, from traced, simulated chains of contour length $L$=300 nm and with different, uniform values of bending stiffness, given by persistence lengths of 30 nm ($N$=73 chains), 85 nm ($N$=78 chains), and 140 nm ($N$=76 chains) for the top, middle and bottom rows, respectively. Three different pixel sizes for the simulated images were tested, with the 4 nm pixel size (right column) representing that used in the experimental scans of this work. Segment lengths of $\Delta s \geq 30$ nm recover the input persistence length within error. Error range is shown as 95% confidence interval.

(α1) SP – KGDCGGSGCGKCDCHGV 44
(α2) SP – LLAQSVLGGVKKLDVPCGGRDCSGGCQCYPEKGARGQPGAVGPQGYNGPPGLQGFPGLQ 84

KGQKGERGLPGLQGVIGFPGMQGPEGPHGPPGQKGDAGEPGLPGTKGTRGPPGAAGYPGNPGLPG 109
GRKGDKGERGVPGPTGPKGDVGARGVSGFPGADGIPGHPGQGGPRGRPGYDGCNGTRGDAGPQGP 149

IPGQDGPPGPPGIPGCNGTKGERGPLGPPGLPGFSGNPGPPGLPGMKGDPGEILGHVPGTLLKGE 174
SGSGGFPGLPGPQGPKGQKGEPYALSKEDRDKYRGEPGEPGLVYQGPPGRPGPIGQMGPMGAPGG 214

RGFPGIPGMPGSPGLPGLQGPVGPPGFTGPPGPPGPPGPPGEKGQMGSSFQGPKGDKGEQGVSGP 239
RPGPPGPPGPKGQPGNRGLGFYGQKGEKGDIGQPGPNGIPSDITLVGPTTSTIHPDLYKGEKGDE 279

PGVPGQAQVKEKGDFAPTGEKGQKGEPGFPGVPGYGEKGEPGKQGPRGKPGKDGEKGERGSPGIP 304
GEQGIPGVISKGEEGIMGFPGIRGFPGLDGEKGVVGQKGSRGLDGFQGPSGPRGPKGERGEQGPP 344

GDSGYPGLPGRQGPQGEKGEAGLPGPPGTVIGTMPLGEKGDRGYPGAPGLRGEPGPKGFPGTPGQ 369
GPSVYSPHPSLAKGARGDPGFQGAHGEPGSRGEPGEPGTAGPPGPSVGDEDSMRGLPGEMGPKGF 409

PGPPGFPTPGQAGAPGFPGERGEKGDQGFPGVSLPGPSGRDGAPGPPGPPGPPGQPGHTNGIVEC 434
SGEPGSPARYLGPPGADGRPGPQGVPGPAGPPGPDGFLFGLKGSEGRVGYPGPSGFPGTRGQKGW 474

QPGPPGDQGPPGTPGQPGLTGEVGQKGQKGESCLACDTEGLRGPPGPQGPPGEIGFPGQPGAKGD 499
KGEAGDCQCGQVIGGLPGLPGPKGFPGVNGELGKKGDQGDPGLHGIPGFPGFKGAPGVAGAPGPK 539

RGLPGRDGLEGLPGPQGSPGLIGQPGAKGEPGEIFFDMRLKGDKGDPGFPGQPGMPGRAGTPGRD 564
GIKGDSRTITTKGERGQPGIPGVHGMKGDDGVPGRDGLDGFPGLPGPPGDGIKGPPGDAGLPGVP 604

GHPGLPGPKGSPGSIGLKGERGPPGGVGFPGSRGDIGPPGPPGVGPIGPVGEKGQAGFPGGPGSP 629
GTKGFPGDIGPPGQGLPGPKGERGFPGDAGLPGPPGFPGPPGPPGTPGQRD**C**DTGVKRPIGGGQQ 669

*Loop 1*

GLPGPKGEAGKVVPLPGPPGAAGLPGSPGFPGPQGDRGFPGTPGRPGIPGEKGAVGQPGIGFPGL 694
VVVQPG**C**IEGPTGSPGQPGPPGPTGAKGVRGMPGFPGASGEQGLKGFPGDPGREGFPGPPGFMGP 734

PGPKGVDGLPGEIGRPGSPGRPGFNGLPGNPGPQGQKGEPGIGLPGLKGQPGLPGIPGTPGEKGS 759
RGSKGTTGLPGPDGPPGPIGLPGPAGPPGDRGIPGEVLGAQPGTRGDAGLPGQPGLKGLPGETGA 799

*Loop 2*

IGGPGVPGEQGLTGPPGLQGIRGDPGPPGVQGPAGPPGVPGIGPPGAMGPPGGQGPPGSSGPPGI 824
PGFRGSQGMPGMPGLKGQPGFPGPSGQPGQSGPPGQHGFPGTPGREGPLGQPGSPGLGGLPGDRG 864

KGEKGFPGFPGLDMPGPKGDKGSQGLPGLTGQSGLPGLPGQQGTPGVPGFPGSKGEMGVMGTPGQ 889
EPGDPGVPGPVGMKGLSGDRGDAGMSGERGHPGSPGFKGMAGMPGIPGQKGDRGSPGMDGFQGML 929

PGSPGPAGTPGLPGEKGDHGLPGSSGPRGDPGFKGDKGDVGLPGMPGSMEHVDMGSMKGQKGDQG 954
GLKGRQGFPGTKGEAGFFGVPGLKGLPGEPGVKGNRGDRGPPGPPPLILPGMKDIKGEKGDEGPM 994

EKGQIGPTGDKGSRGDPGTPGVPGKDGQAGHPGQPGPKGDPGLSGTPGSPGLPGPKGSVGGMGLP 1019
GLKGYLGLKGIQGMPGVPGVSGFPGLPGRPGFIKGVKGDIGVPGTPGLPGFPGVSGPPGITGFPG 1059

GSPGEKGVPGIPGSQGVPGSPGEKGAKGEKGQSGLPGIGIPGRPGDKGDQGLAGFPGSPGEKGEK 1084
FTGSRGEKGTPGVAGVFGETGPTGDFGDIGDTVDLPGSPGLKGERGITGIPGLKGFFGEKGAAGD 1124

```
GSAGTPGMPGSPGPRGSPGNIGHPGSPGLPGEKGDKGLPGLDGVPGVKGEAGLPGTPGPTGPAGQ  1149
IGFPGITGMAGAQGSPGLKGQTGFPGLTGLQGPQGEPGRIGIPGDKGDFGWPGVPGLPGFPGIRG  1189

KGEPGSDGIPGSAGEKGEQGVPGRGFPGFPGSKGDKGSKGEVGFPGLAGSPGIPGVKGEQGFMGP  1214
ISGLHGLPGTKGFPGSPGVDAHGDPGFPGPTGDRGDRGEANTLPGPVGVPGQKGERGTPGERGPA  1254

PGPQGQPGLPGTPGHPVEGPKGDRGPQGQPGLPGHPGPMGPPGFPGINGPKGDKGNQGWPGAPGV  1279
GSPGLQGFPGISPPSNISGSPGDVGAPGIFGLQGYQGPPGPPGPNALPGIKGDEGSSGAAGFPGQ  1319

PGPKGDPGFQGMPGIGGSPGITGSKGDMGLPGVPGFQGQKGLPGLQGVKGDQGDQGVPGPKGLQG  1344
KGWVGDPGPQGQPGVLGLPGEKGPKGEQGFMGNTGPSGAVGDRGPKGPKGDQGFPGAPGSMGSPG  1384

PPGPPGPYDVIKGEPGLPGPEGPPGLKGLQGPPGPKGQQGVTGSVGLPGPPGVPGFDGAPGQKGE  1409
IPGIPQKIAVQPGTLGPQGRRGLPGALGEIGPQGPPGDPGFRGAPGKAGPQGRGGVSAVPGFRGD  1449

TGPFGPPGPRGFPGPPGPDGLPGSMGPPGTPSVDH 1444...NC1
QGPMGHQGPVGQEGEPGRPGSPGLPGMPGRSVSIG 1484...NC1

               Aligned from here <--'
```

**Figure S5. Amino acid sequences of the collagen IV collagenous domain.** The α1 (P02463, in blue) and α2 (P08122, in red) amino acid sequences were aligned at the first Gly-X-Y overlap from the NC1 domain, as the assembly of collagen is initiated at that end. The signal peptide sequences for both chains are not displayed in this alignment, but are denoted by SP at the start of the sequence. The underlined portions at the beginning of the sequences correspond to the 7S-forming regions of collagen IV. The highlighted segments are interruptions in the triple-helical-defining $(Gly\text{-}X\text{-}Y)_nG$ amino acid sequence. The triple-helical regions have been defined as $(Gly\text{-}X\text{-}Y)_nG$ with $n \geq 4$. The longest interruption, 26 amino acids long in α2, has two cysteines (bolded in black) near its edges, and is treated as loop 1 in the "two loops" sequence alignments. A second interruption in α2 ("loop 2"; centered at LGA) is also removed from the backbone contour in the two loops alignments.

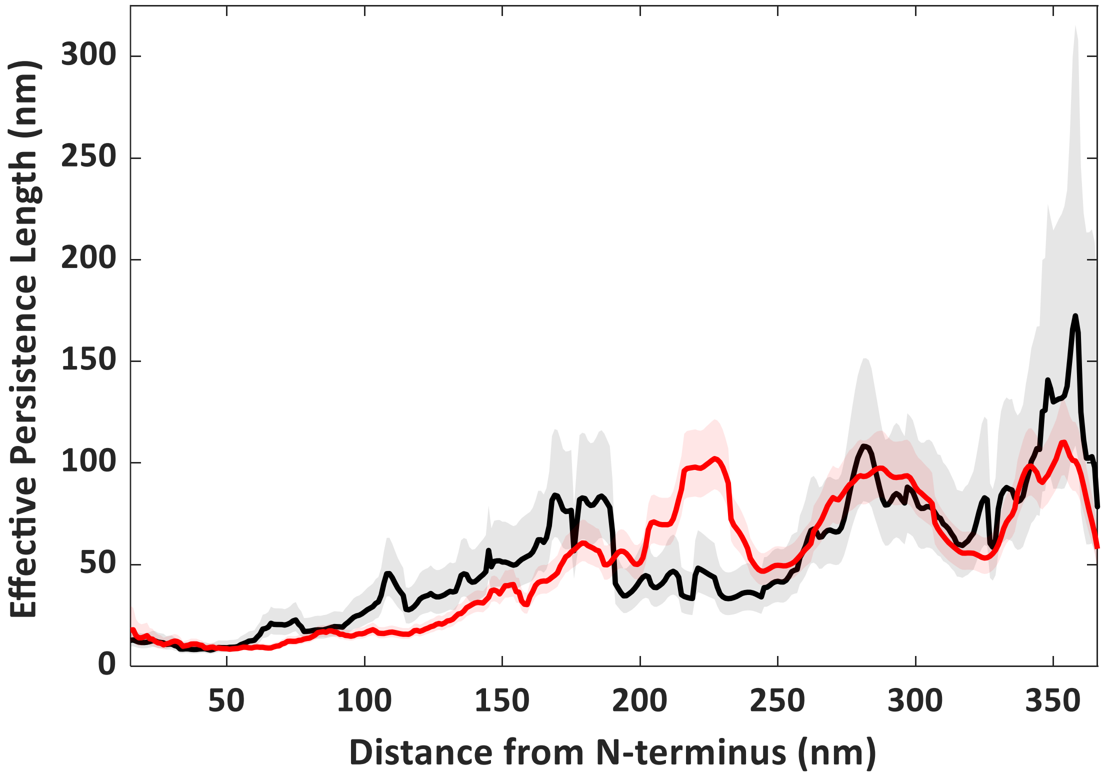


**Figure S6. Flexibility profiles of collagen IV traced in both directions.** Collagen type IV was deposited from 100 mM KCl, 1 mM HCl and traced in both directions. The shaded curves represent 95% confidence intervals on the effective persistence length estimate $p(s; \Delta s)$. The profile in black was traced from the N terminus (7S domain) towards the C terminus (NC1 domain) and was calculated from $N$ = 84 chains. The profile in red (reproduced from Fig. 4) was traced from the C terminus (NC1 domain) towards the N terminus (7S domain) and was calculated from $N$ = 262 chains. The two profiles displayed are aligned at the N terminus and appear comparable, with the exception of the region around 225 nm from the N terminus. This may be a result of the many fewer chains that were traced from the N terminus.

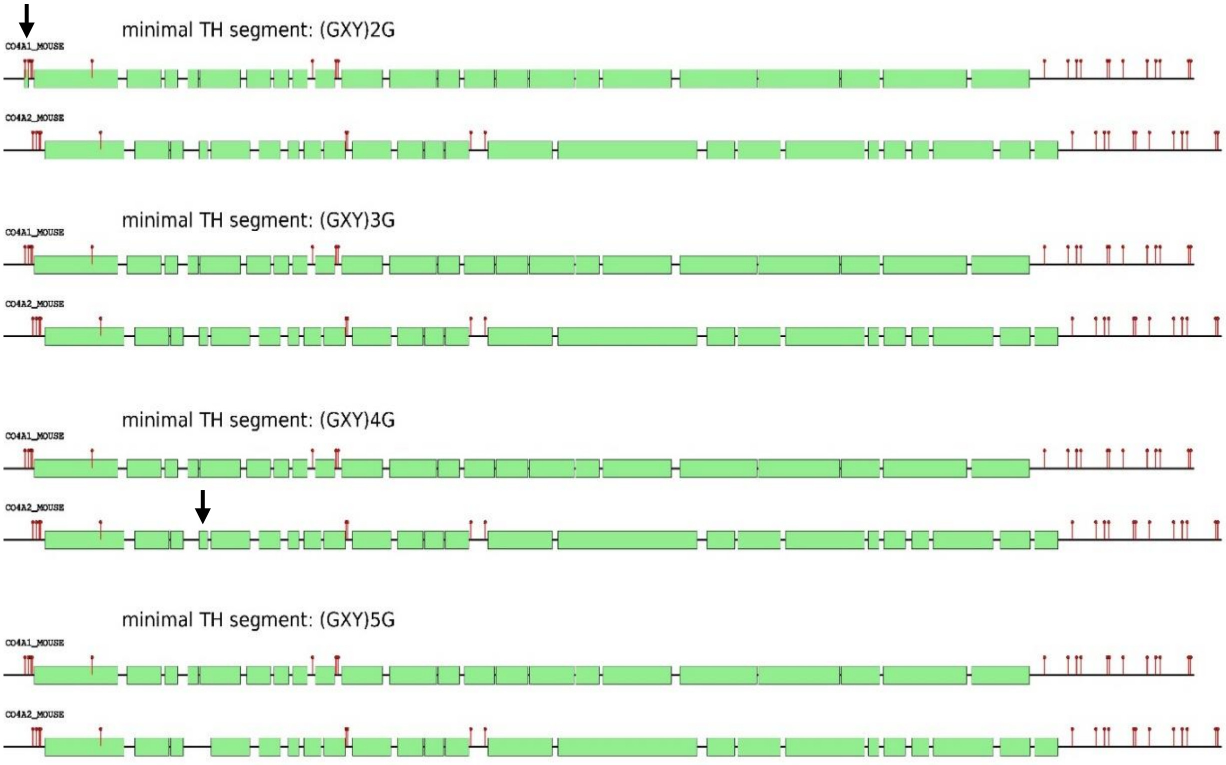


**Figure S7. Varying assumptions of the minimum number of tripeptide units required to form triple-helical segments.** Schematic depiction of sequences when varying the minimal sequence requirements for a triple helix from $(\text{Gly-X-Y})_n\text{G}$, $n$=2-5. Each of the two bold arrows indicates a triple-helix-forming segment that is lost when increasing $n$ (from 2 to 3, and from 4 to 5). All other triple-helical segments are longer than $(\text{GXY})_4\text{G}$.

### Supporting Text 2

#### Variable flexibility model fitting

The variable flexibility model was first used to define a position-dependent profile $[p_{\mathrm{in,i}}]=[p_{\mathrm{in,1}}, p_{\mathrm{in,2}}, \ldots, p_{\mathrm{in,N}}]$ in which the input flexibility (via local persistence length) is defined at each position $i=(1, 2, \ldots, N)$ along the chain backbone. For the simulated chains, each $p_{\mathrm{in,i}}$ can take the value $p_{\mathrm{r}}$ or $p_{\mathrm{f}}$, corresponding to a rigid or flexible monomer in the chain, while for collagen IV, each $p_{\mathrm{in,i}}$ can take one of four values: $p_0$, $p_1$, $p_2$ and $p_3$ (Table 1 in main text) The profile $[p_{\mathrm{in,i}}]$ was fit to the measured persistence length profile as follows, with the aim of determining values for the local flexibilities ($p_{\mathrm{r}}$ and $p_{\mathrm{f}}$; or $p_0$, $p_1$, $p_2$ and $p_3$).

1. A `filter` window of length $\Delta s$ was used to determine the effective persistence length profile, which depends on all local persistence lengths within this window as given by equation (S6). For simulated chains, whose monomer spacing is defined in nanometers, `filter` = $\Delta s$. For collagen IV chains, the monomer spacing is defined in amino acids, and so a scaling factor `nmaa` (nanometers / amino acid) is applied to express the filter length as the number of amino acid steps:
$$\text{filter} = \frac{\Delta s}{\text{nmaa}}. \tag{S15}$$
The effective persistence length profile $[p_{\mathrm{eff},i}]$ is then given by
$$p_{\mathrm{eff},i} = \frac{\text{filter}}{\sum_{i}^{i+\text{filter}}\left(p_{\mathrm{in},j}{}^{-1}\right)}, \tag{S16}$$
where the step spacing remains in its original units (e.g. amino acid steps for collagen IV). The $p_{\mathrm{eff}}$ profiles are shorter than the initial profiles: $i$=(1:$N$-`filter`+1). In the results reported here, $\Delta s$ = 30 nm for all measurements and fitting.
2. A `stagger` parameter was used, which represents the standard deviation of chain starting positions in the traced population. This was incorporated to account for variability in identifying the starting position of the chains from images. `stagger` is defined in units of nanometers and converted, for collagen chains, to amino acid steps as for the filter window length (S15). A Gaussian smoothing window was applied to the profile from (S16), with contributions from positions up to ±3 standard deviations included:
$$p_{\mathrm{eff,stag},i} = \sum_{i}^{i+6\times\text{stagger}} P(j)\,p_{\mathrm{eff},j}. \tag{S17}$$
Here
$$P(j) = Ae^{-\frac{\left(j-(i+3\times\text{stagger})\right)^2}{2(\text{stagger})^2}} \tag{S18}$$
is the Gaussian smoothing function, where $A$ is a normalization constant determined so that $\sum_{i}^{i+6\times\text{stagger}} P(j) = 1$. The $p_{\mathrm{eff,stag}}$ profiles run from $i$=(1:$N$-`filter`-6×`stagger`+1).
3. Prior to fitting, the collagen chain $p_{\mathrm{eff,stag}}$ profiles in amino acid steps were converted into nanometer integer increments, via linear interpolation. The profiles then run from s = ((`filter`/2 + (6*`stagger`-1)/2) : (`length` - `filter`/2 - 6*`stagger`/2 + 1/2)).

4. An `offset` parameter was added to the contour positions ($s \rightarrow s$ + `offset`), which has the effect of linearly displacing the simulated profile relative to the measured profile. `offset` accounts for systematic errors in determining the start position of the chain (for example, if it is obscured by the NC1 domain).
5. Before fitting, the simulated and traced chain persistence length profiles must be the same length. Non-overlapping beginning and/or end segments of $p_{\text{eff,stag}}(s)$ and/or $p(s)$ were removed to generate two linear arrays of identical length and range in $s$ (and which have identical increments along the contour, of 1 nm).
6. The difference between $p_{\text{eff,stag}}(s)$ and $p(s)$ was minimized by varying $p_0$, $p_1$, $p_2$ and $p_3$ [$p_r$ and $p_f$ in the case of simulated chains], and weighting the estimates of each value of $p(s)$ in the traced chain profile by its variance (estimated from equation (S13) by assuming normally distributed errors):
$$f(p_0, p_1, p_2, p_3) = \sum_s \frac{\left[p(s) - p_{\text{eff,stag}}(s)\right]^2}{\text{Var}(s)}. \quad \text{(S19)}$$
Persistence length values ($p_0$, $p_1$, $p_2$ and $p_3$) were constrained to lie within the range [0,200] nm. The $\chi_r^2$ value that resulted from minimizing the function $f$ with the best-fit values of $p_0$, $p_1$, $p_2$ and $p_3$ was recorded:
$$\chi_r{}^2 = \frac{f_{\text{min}}(p_0, p_1, p_2, p_3)}{N_{\text{pts}} - N_{\text{params}}}. \quad \text{(S20)}$$
$N_{\text{pts}}$ is the length of the resulting $p(s)$ and $p_{\text{eff,stag}}(s)$ arrays used for minimizing $f$, and $N_{\text{params}} = 4$ for collagen and 2 for the simulated chains.
7. Steps 2-6 were repeated for different parameters `nmaa`, `stagger` and/or `offset` to determine the values of $p_0$, $p_1$, $p_2$ and $p_3$ that minimized $\chi_r^2$. In practice, this was implemented in a series of nested loops, with `nmaa` taking possible values of [0.27, 0.29, 0.31, 0.33, 0.35] nm/aa, and `stagger` and `offset` each taking integer values. For simulated chains, an optimal `stagger` = 5 nm was determined. For collagen IV, we found $\chi_r^2$ to decrease as `stagger` was increased, in effect smearing out and making more homogeneous the simulated chain $p_{\text{eff,stag}}$ profile. Thus, we kept `stagger` = 5 nm fixed, while varying `offset` and `nmaa` to determine their optimal values. These parameters, and the resulting best-fit values of $p_0$, $p_1$, $p_2$ and $p_3$, are presented in Table S1 for some of the tested chain alignments.

All data fitting was implemented within MATLAB (4).

Obtaining a well resolved effective persistence length profile from experimental images relies on accurate contour tracing and chain start-point determination. Experimentally, AFM images of collagen IV were obtained with settings that saturated the intensity in the NC1 domain. This led to a plateau-like intensity profile of the NC1 domains (typical diameter ~12 nm), from which we estimate the error of edge determination to be <2 pixels. This is commensurate with the 5 nm value of `stagger` used in smoothing the model flexibility profiles. Further refining this start-point of chain tracing would decrease blurring of chain registry and improve the mapping of the underlying flexibility profile.

**Table S1: Model outputs of flexibility for different chain alignments.** The model was fit to the $p(s)$ profile obtained for 100 mM KCl, 1 mM HCl using a $\Delta s$ = 30 nm filter window. All fits imposed a $\sigma$ = 5 nm chain stagger and constrained the values of each persistence length to $0 \leq p_i \leq$ 200 nm and offset to $-20 \leq offset \leq 0$. Red values highlight those results that were optimized at the fitting boundary and thus did not minimize within the parameter range. The fits for bolded alignments are shown in the main manuscript. Clustal alignments vary in the loops imposed and assumptions about their contributions to the main contour. Two loops alignment names indicate the number of amino acids looped out from the α2 chain around residues 656-676 (disulfide-bridged loop) and 771-774 (Fig. S6) (5). Detailed sequence alignments are provided in a supporting file. The last column indicates whether α1 441D and α2 456R are aligned, as found for this integrin-binding region in the human homolog (6). Only the final, italicized alignment includes staggered starting positions in the three chains; here, the relative chain offset produces the local register found for the mapped integrin-binding site in human type IV collagen (6).

| Chain Alignment | Offset (nm) | nm/aa | $p$ (nm) | $p_1$ (nm) | $p_2$ (nm) | $p_3$ (nm) | $\chi^2_r$ | DDR aligned? |
|---|---|---|---|---|---|---|---|---|
| Linear | -11 | 0.33 | 154 | 10.8 | 2.3 | 200 | 7.7 | No |
| Clustal A | -17 | 0.29 | 146 | 200 | 15 | 1.9 | 12.2 | Yes |
| **Clustal B** | -17 | 0.29 | 109 | 86 | 42 | 2.0 | 12.8 | Yes |
| Clustal C | -20 | 0.31 | 125 | 33 | 11 | 1.3 | 14.2 | Yes |
| One loop | -7 | 0.33 | 154 | 9.9 | 1.8 | 200 | 10.6 | No |
| Two loops (19,3) | -11 | 0.29 | 163 | 20 | 6.6 | 1.7 | 14.4 | No |
| Two loops (19,4) | -10 | 0.29 | 138 | 28 | 8.1 | 1.8 | 14.0 | No |
| **Two loops (21,4)** | -10 | 0.29 | 105 | 83 | 21 | 1.9 | 12.6 | Yes |
| Two loops (21,5) | -9 | 0.29 | 102 | 112 | 24 | 2.0 | 12.8 | No |
| *Two loops (21,4) – staggered* | *-7* | *0.29* | *129* | *200* | *3.5* | *2.1* | *14.0* | *Yes* |

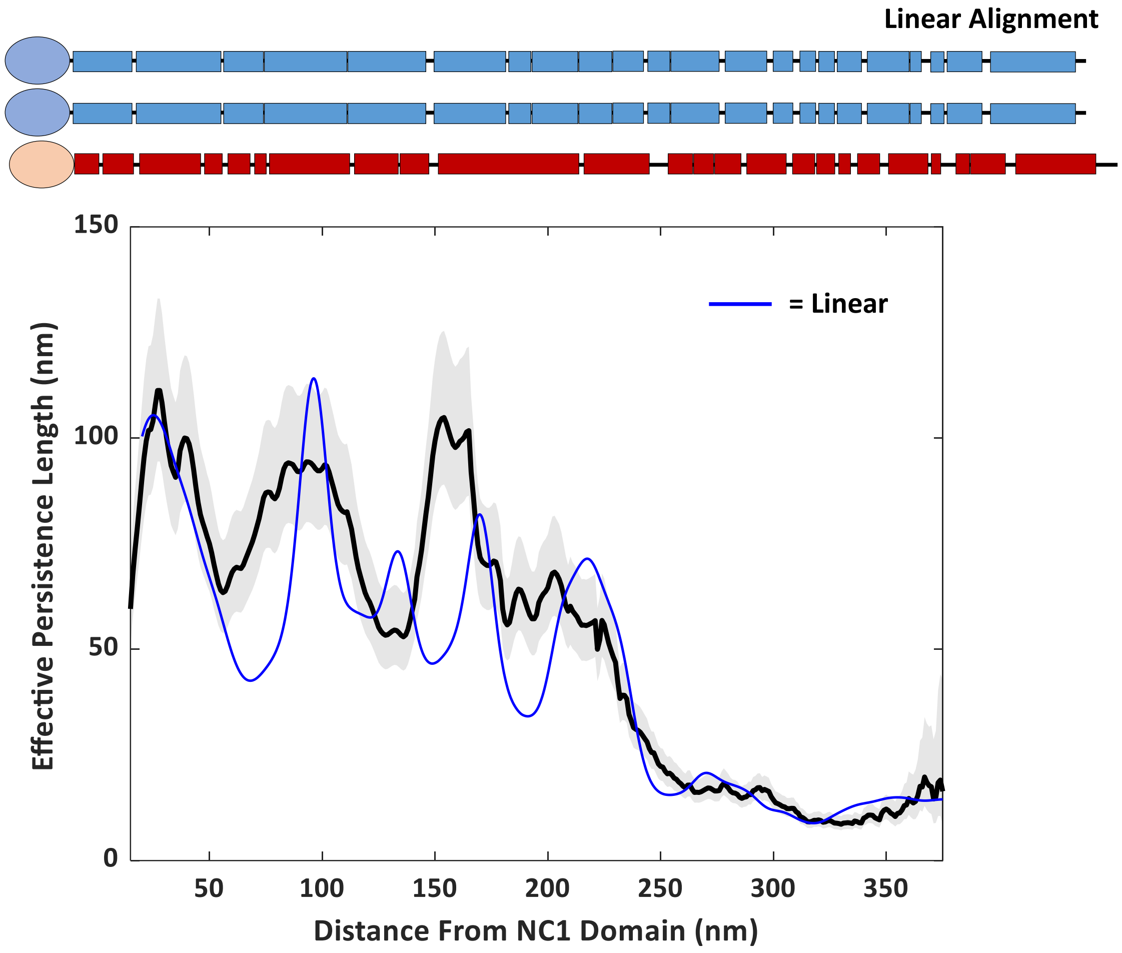


**Figure S8. Four-class flexibility model using a linear chain alignment**. The persistence length profile of collagen IV deposited from 100 mM KCl 1 mM HCl is aligned with the amino acid sequence representations using the offset (-11 nm) and 0.33 nm/aa conversion parameters obtained from the fitting procedure. The model produced for an overlapping interaction $p_3$ = 200 nm, the maximum value allowed for fitting, an unphysical value. However, fitting $p^*(s)$ by using this linear chain alignment did capture the rigidity in the collagenous region adjacent to the NC1 domain.

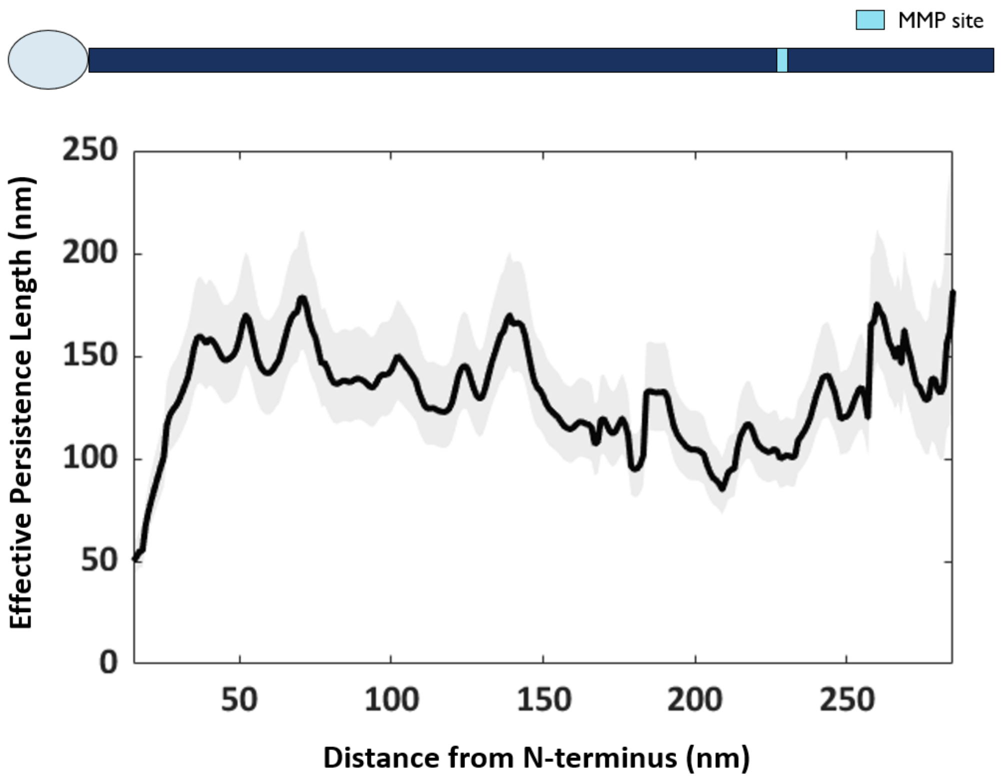


**Figure S9. Position-dependent flexibility profile of collagen pN-III.** Position-dependent persistence length map of collagen pN-III traced from the N-terminus. The profile is aligned with an amino acid sequence representation where the MMP site location is marked. Shaded curves represent 95% confidence intervals on the effective persistence length estimate $p(s; \Delta s)$. The profile was calculated from $N$ = 267 chains.

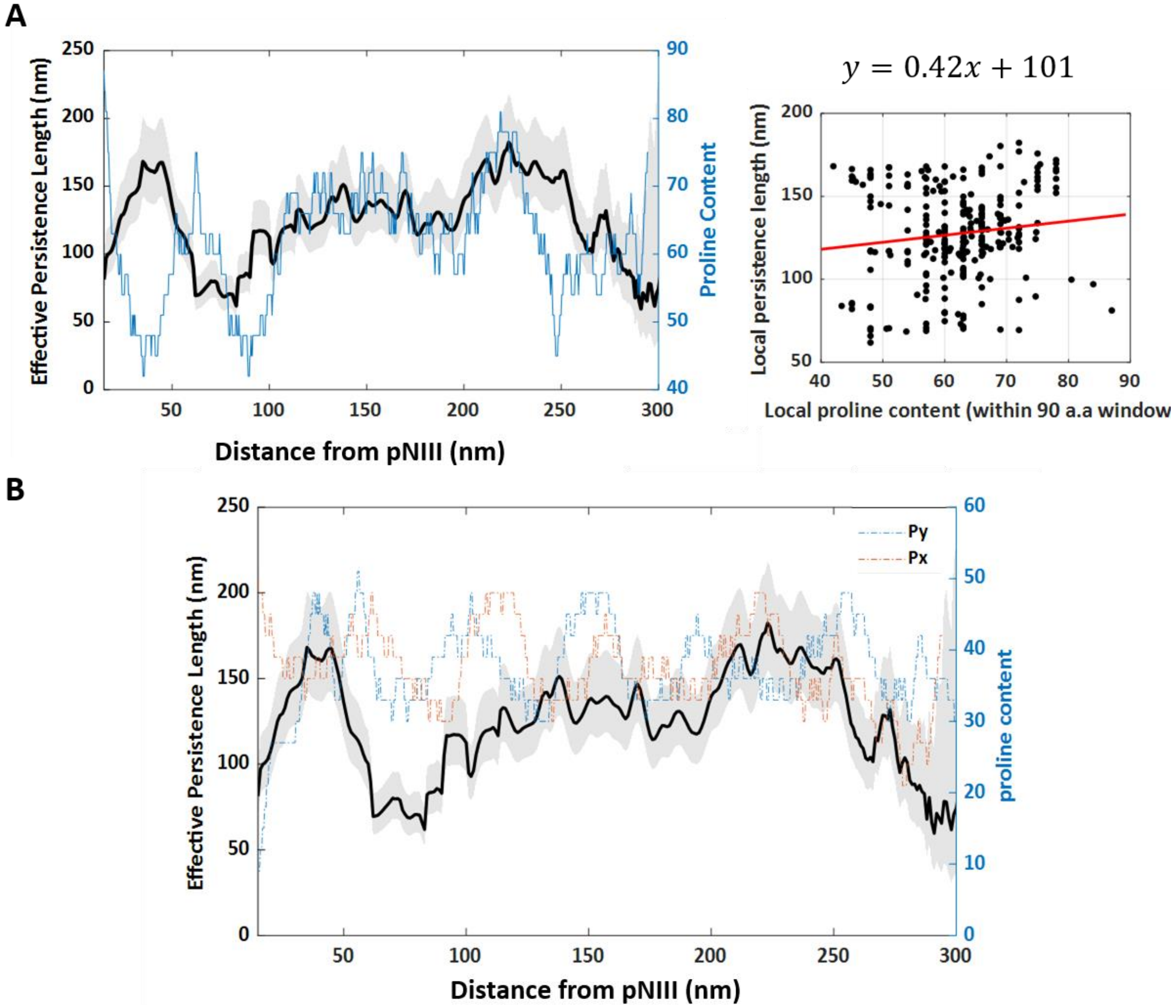


**Figure S10. Local imino-acid (proline) content and persistence length profile of pN-III collagen.** (A) The local proline content (calculated over a 90-amino-acid sliding window within each of the three chains) and effective persistence length of pN-III collagen (Q08E14) are uncorrelated, shown by a linear correlation coefficient of $R^2 = 0.017$. (B) There appears to be no strong correlation between either X- or Y-positioned proline content and effective persistence length. X-positioned prolines are expected to be unmodified, while Y-positioned prolines are expected to be 4-hydroxylated. Proline content in bovine pN-III collagen is given by the number of proline residues found in three α1(III) (Q08E14) chains, within a 90-amino-acid sliding window centered at the position noted. Proline content was determined at positions centered every 3 amino acids along the chain and was linearly interpolated to obtain values at nanometer spacing.

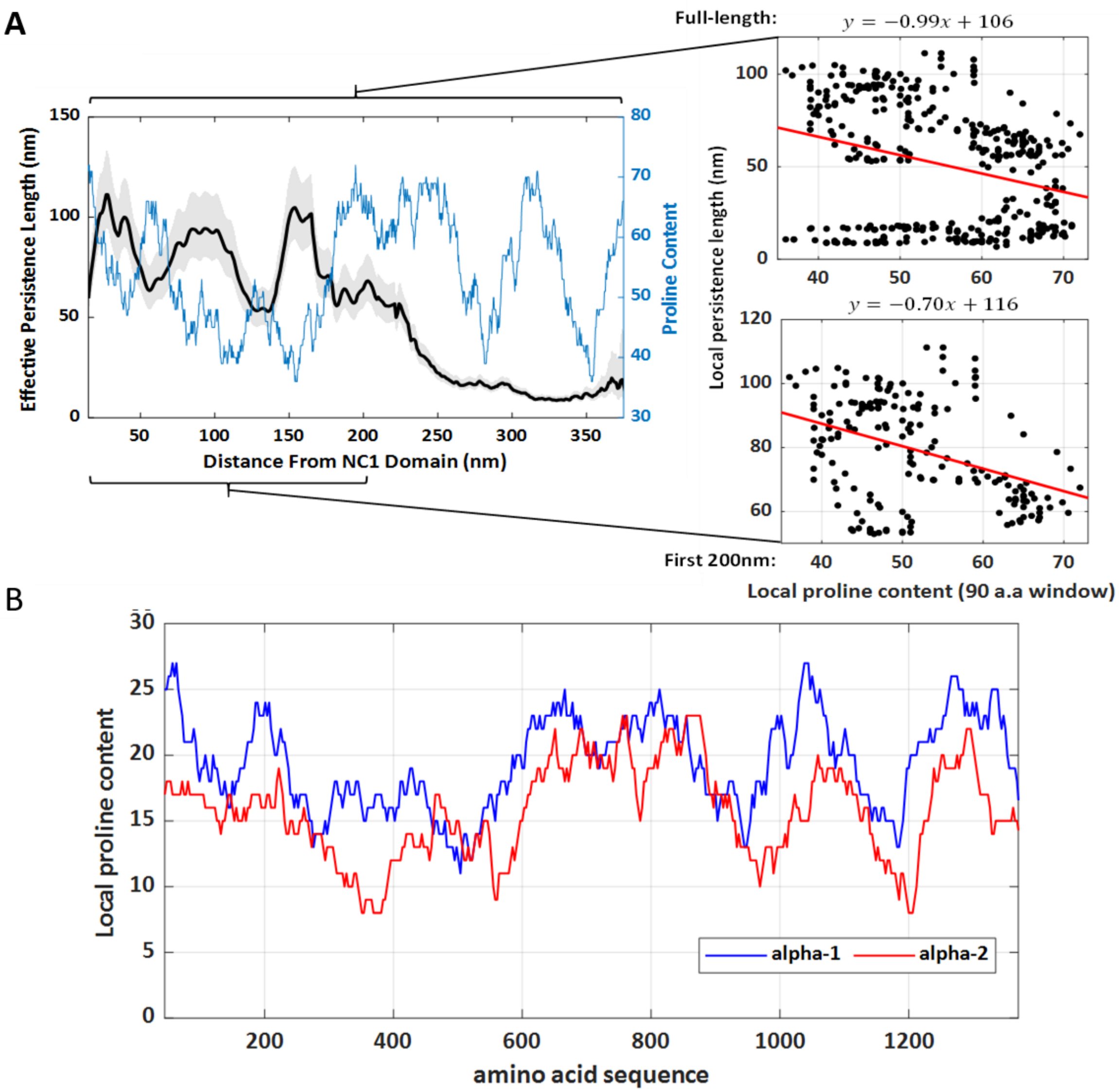


**Figure S11. Local imino-acid (proline) content and persistence length profile of collagen IV.** (A) The local proline content and effective persistence length appear anti-correlated in the C-terminal half of collagen IV, and otherwise uncorrelated. Quantitative analysis, however, reveals no statistical correlation between proline content and flexibility, neither for the full length of the chain ($R^2$ = 0.084) nor for the first 200 nm from the NC1 domain ($R^2$ = 0.15). The analysis assumes the three chains of collagen IV to be linearly aligned (no loops) and uses a conversion of 0.29 nm / amino acid. (C) Proline content in mouse collagen IV is given by the number of proline residues found in α1(IV) (P02463) and α2(IV) (P08122) chains, within a 90-amino-acid sliding window centered at the position noted. Proline content was determined at positions centered every 3 amino acids along the chain and was linearly interpolated to obtain values at nanometer spacing.

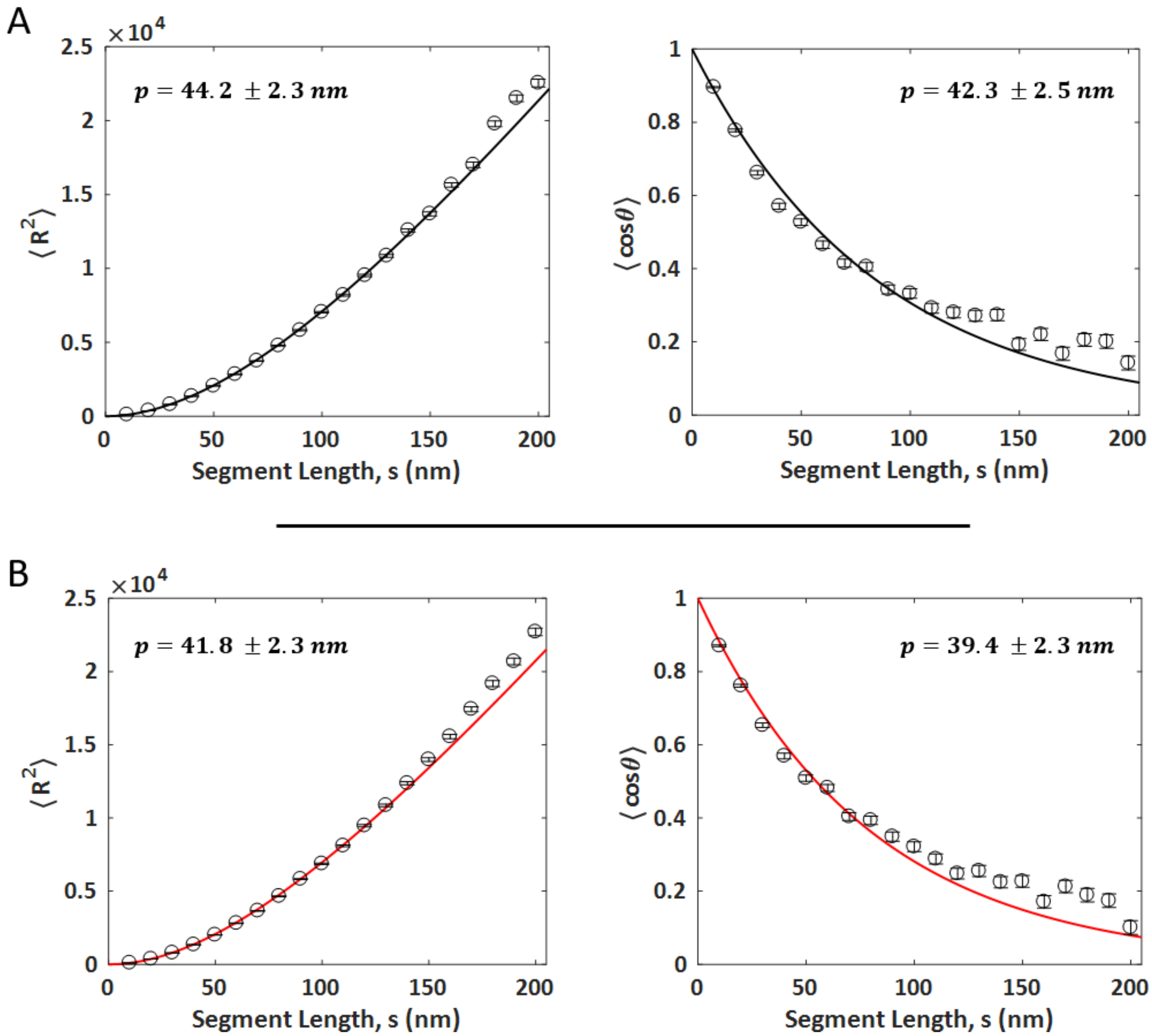


**Figure S12. Homogeneous worm-like chain determination of persistence lengths of collagen IV in different solution conditions.** The persistence lengths were obtained using $\langle R^2(\Delta s)\rangle$ (plots to the left) and $\langle \cos\theta(\Delta s)\rangle$ (plots to the right) analyses. The data correspond to collagen type IV deposited from A) TBS (chloride-containing), and from B) sodium acetate buffer.

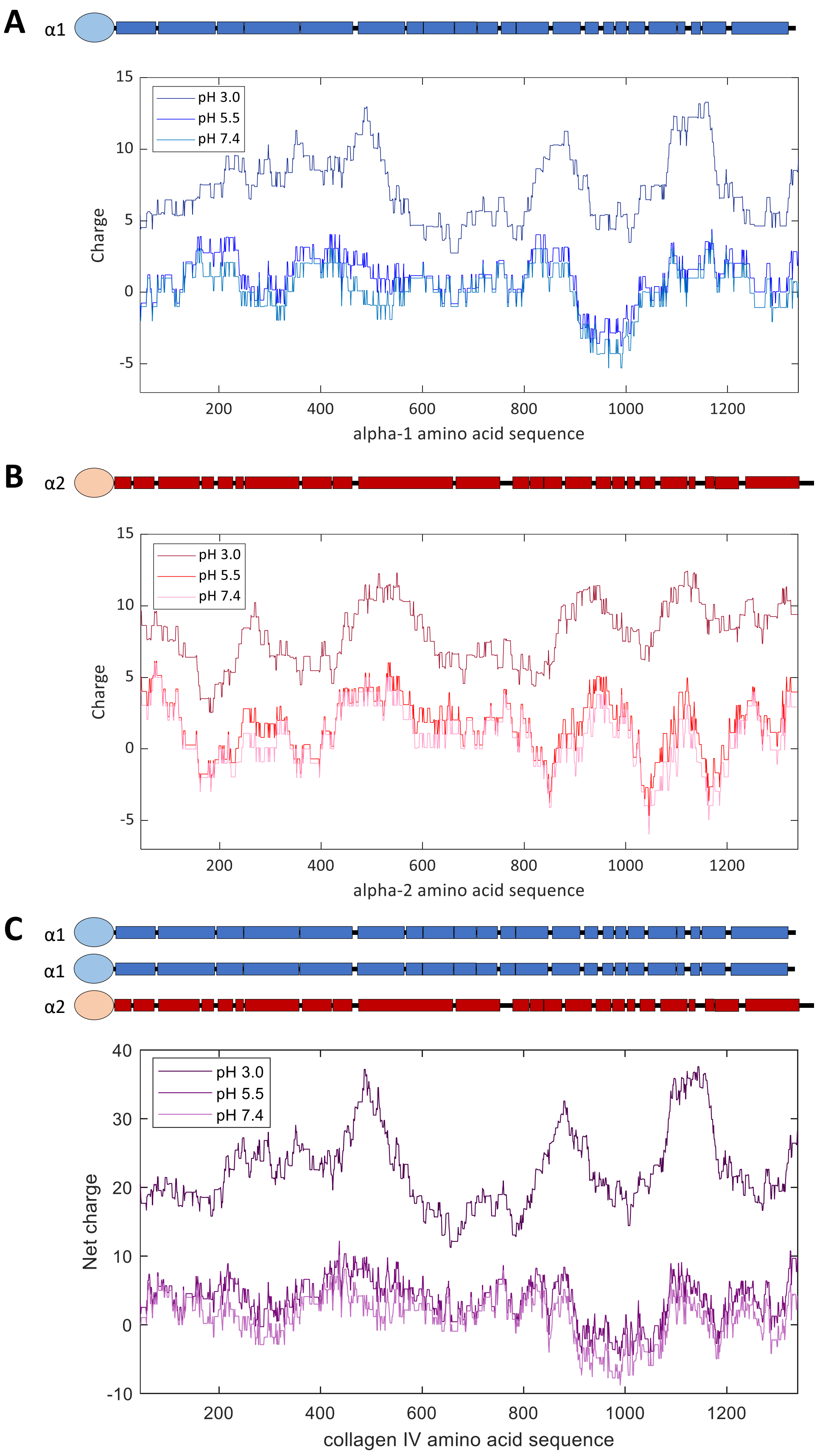

A
α1
pH 3.0
pH 5.5
pH 7.4
Charge
15
10
5
0
-5
200
400
600
800
1000
1200
alpha-1 amino acid sequence
B
α2
pH 3.0
pH 5.5
pH 7.4
Charge
15
10
5
0
-5
200
400
600
800
1000
1200
alpha-2 amino acid sequence
C
α1
α1
α2
pH 3.0
pH 5.5
pH 7.4
Net charge
40
30
20
10
0
-10
200
400
600
800
1000
1200
collagen IV amino acid sequence

**Figure S13. Estimated charge profiles of the α1 and α2 chains of collagen IV at different pH.** The charge on each amino acid at each pH was calculated using the Henderson-Hasselbach equation and the pKa of the amino acid side chain. Amino acids considered in the calculation, with their corresponding assumed pKa, are Aspartate, $pKa_D$ = 3.9; Glutamate, $pKa_E$ = 4.3; Histidine, $pKa_H$ = 6.1; Cysteine, $pKa_C$ = 8.3; Tryptophan $pKa_Y$ = 10.1; Lysine, $pKa_K$ = 10.5; Arginine, $pKa_R$ = 12.0. The local charge along the sequence was averaged over 90 amino acids centred at each amino acid along the sequence. This estimate assumes that the pKa and charge on each amino acid are unaffected by the surrounding amino acids. **(A)** Charge profile for α1(IV) (P02463). **(B)** Charge profile for α2(IV) (P08122). **(C)** Net charge profile of collagen IV, assuming the three chains of collagen IV to be linearly aligned (no looping).

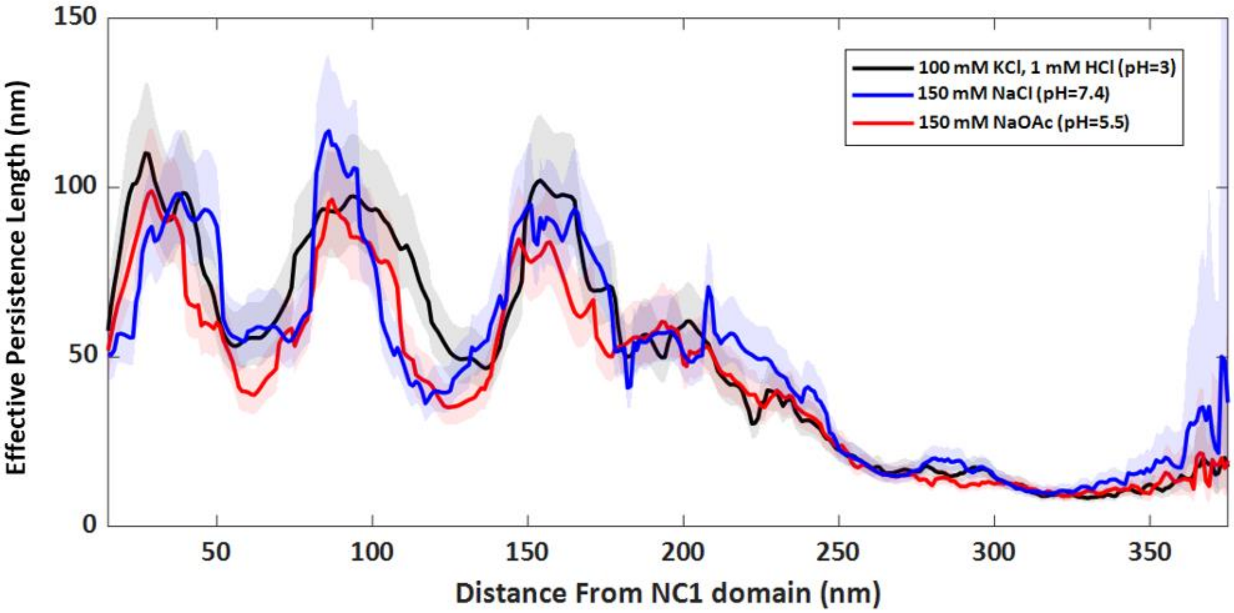


**Figure S14. Superposition of experimental persistence length profiles.** The position-dependent effective persistence length profile of collagen IV changes remarkably little over a pH range from 3 to 7.4, and in the presence or absence of ~150 mM $Cl^-$ ions.

**Supporting References**


1. Landau, L. D., L. P. Pitaevskii, A. M. Kosevich, and E.M.Lifshitz (1986) Theory of Elasticity. Butterworth-Heinemann.

2. Krishnamoorthy,K. (2016) Handbook of statistical distribution with applications (2nd edition). CRC Press: Boca Raton, Florida.

3. Gelman,A. (2013) Bayesian Data Analysis (3rd edition). CRC Press: Boca Raton, Florida.

4. MATLAB Release 2020a. The MathWorks, Inc., Natick, Massachusetts, United States.

5. Kühn,K. (1995) Basement membrane (type IV) collagen. *Matrix Biol.*, **14**, 439–445. https://doi.org/10.1016/0945-053X(95)90001-2

6. Golbik,R., Eble,J.A., Ries,A. and Kühn,K. (2000) The spatial orientation of the essential amino acid residues arginine and aspartate within the α1β1 integrin recognition site of collagen IV has been resolved using fluorescence resonance energy transfer. *J. Mol. Biol.*, **297**, 501–509.
https://doi.org/10.1006/jmbi.2000.3575
